# Reversible inhibition of GluN2B-containing NMDA receptors with an *in situ* red-shifted, photoswitchable antagonist

**DOI:** 10.1101/2023.10.16.562518

**Authors:** Chloé Geoffroy, Romain Berraud-Pache, Nicolas Chéron, Isabelle McCort-Tranchepain, Pierre Paoletti, Laetitia Mony

## Abstract

NMDA receptors (NMDARs) are glutamate-gated ion channels playing a central role in synaptic transmission and plasticity. Dysregulation of NMDARs is linked to various neuropsychiatric disorders, emphasizing the need to understand the functional roles of individual NMDAR subtypes in the brain. GluN2B-containing NMDARs (GluN2B-NMDARs) are particularly important due to both pro-cognitive and pro-excitotoxic roles, although these functions remain under debate. Traditional pharmacological and genetic approaches have important shortcomings in terms of specificity and spatio-temporal resolution, limiting their use in native tissues. We therefore turned to optopharmacology, a technique based on the use of photosensitive ligands, whose activity can be reversibly tuned via illumination with different wavelengths. We developed OptoNAM-3, an azobenzene-based, photoswitchable negative allosteric modulator selective for GluN2B-NMDARs. OptoNAM-3 is a potent inhibitor of GluN2B-NMDARs in its *trans* configuration and inactive in its *cis* configuration. When bound to GluN2B-NMDARs, OptoNAM-3 displays remarkable red-shifting of its photoswitching properties that we attributed to geometric constraints imposed by the binding site onto the ligand azobenzene moiety. OptoNAM-3 allowed fast and reversible photomodulation of GluN2B-NMDAR activity *in vitro* using either UV/green or blue/green light illumination cycles. OptoNAM-3 furthermore acted as a reversible, red-shifted *in vivo* photomodulator of Xenopus tadpole locomotion. By enabling fast and reversible photocontrol of endogenous GluN2B-NMDARs with *in vivo* compatible photochemical properties, OptoNAM-3 should advance our understanding of the role of this class of NMDARs in brain function and dysfunction.

**Significance statement:** This article presents the development and characterization of a photoswitchable negative allosteric modulator (NAM) targeting GluN2B-containing NMDA receptors (GluN2B-NMDARs). Traditional GluN2B-selective NAMs suffer from slow kinetics and irreversible effects, limiting their use in native tissues. OptoNAM-3 emerged as a potent and selective inhibitor of GluN2B-NMDARs, exhibiting fast temporal resolution of action and reversibility both *in vitro* and *in vivo*. OptoNAM-3 furthermore exhibited different spectral properties when in solution or bound to its target, thus behaving as an *in situ* “red-shifted” photodependent antagonist with improved *in vivo* compatibility. This study therefore provides a valuable photoswitchable tool for precise control of NMDAR activity in native tissues. It furthermore reveals the importance of the protein environment on the spectral properties of photosensitive molecules.

## Introduction

Neuronal plasticity, the brain’s ability to continually adapt to its environment or experiences, hinges on the dynamics of chemical synapses. At these specialized neuronal sites, neurotransmitters released from a presynaptic neuron cross the synaptic cleft and activate receptors on the postsynaptic neuron, hence mediating transmission of information from one neuron to another. NMDA receptors (NMDARs) are a class of ionotropic receptors activated by glutamate, the main excitatory neurotransmitter of the Vertebrate central nervous system. They play a central role in synaptic transmission and plasticity but their dysfunction is also involved in many pathologies (1–3). NMDARs are tetramers composed of two GluN1 and two GluN2 (or GluN3) subunits. Each tetramer can either incorporate two identical GluN2 (or GluN3) subunits (di-heteromers) or different GluN2 (or GluN3) subunits (tri-heteromers), each GluN2 subunit conferring to the receptor distinct biophysical and pharmacological properties, as well as different expression and signaling profiles (1, 2). Understanding the functional role of NMDAR individual subtypes in the brain is fundamental to develop new strategies to counteract the deleterious effects of NMDAR deregulation.

Over-activation of NMDARs, as occurring during traumatic brain injury or stroke, induces an excessive increase in intracellular calcium, a process leading to neuronal death (excitotoxicity) (4). This excitotoxicity phenomenon is also observed in neurodegenerative diseases like Alzheimer’s and Parkinson’s diseases (1, 2, 5–8). NMDAR overactivation furthermore occurs in other pathologies such as epilepsy, neuropathic pain and depression (1, 2, 5, 9, 10). Multiple studies point to a specific role of GluN2B-NMDARs in triggering excitotoxicity, although this role is debated (1, 5, 8, 11). To counteract the deleterious effects of GluN2B-NMDAR overactivation, a large number of negative allosteric modulators (NAMs) specific for GluN2B-NMDARs were developed in the late 90s – early 2000s (5, 12–15). These antagonists, of which ifenprodil is the lead compound (16–18), displayed neuroprotective properties *in vitro* and *in vivo* with reduced adverse effects compared to broad spectrum antagonists (1, 5, 14, 19–21). So far, however, all of these compounds failed in clinical trials because of a lack of effect or a narrow therapeutic window (5, 21, 22).

Ifenprodil derivatives bind at the level of the NMDAR N-terminal domains (NTDs) (23, 24), bilobar domains preceding the agonist-binding domain (ABD) and that constitute a hub for allosteric modulation in NMDARs (2, 3, 5, 12, 25) (see Fig. 2A for NMDAR subunit architecture). At this level, these compounds induce their inhibition by interacting with the upper lobe of GluN1 NTD and with the upper and lower lobes of the GluN2B NTD (23, 24, 26), which favors entry of NMDARs into an inhibited state (26–30). Some of these GluN2B-specific NAMs, such as ifenprodil, Ro25-6981 (31) or CP-101,606 (32, 33) are currently used as standard pharmacological tools to specifically target GluN2B-NMDARs in native tissues and have proven useful to investigate the contributions of this receptor subtype to several physiological and pathological processes. However, use of these compounds in native tissues faces serious limitations due to their slow association and dissociation kinetics. In recombinant systems, time constants of association of GluN2B-specific NAMs are indeed in the seconds to tens of seconds time-range at saturating compound concentration, while time constants of dissociation are in the tens of seconds to minute time-range (34–37). These slow kinetics are even more marked in native tissues. In brain slices, for instance, GluN2B-specific NAMs take tens of minutes to elicit their inhibitory effect, which is then irreversible (see, for instance, refs (38, 39)). It is thus important to develop GluN2B-selective inhibitors with improved temporal resolution and reversibility of action for a dynamic control of GluN2B-NMDARs in native tissues.

**Fig. 1.**
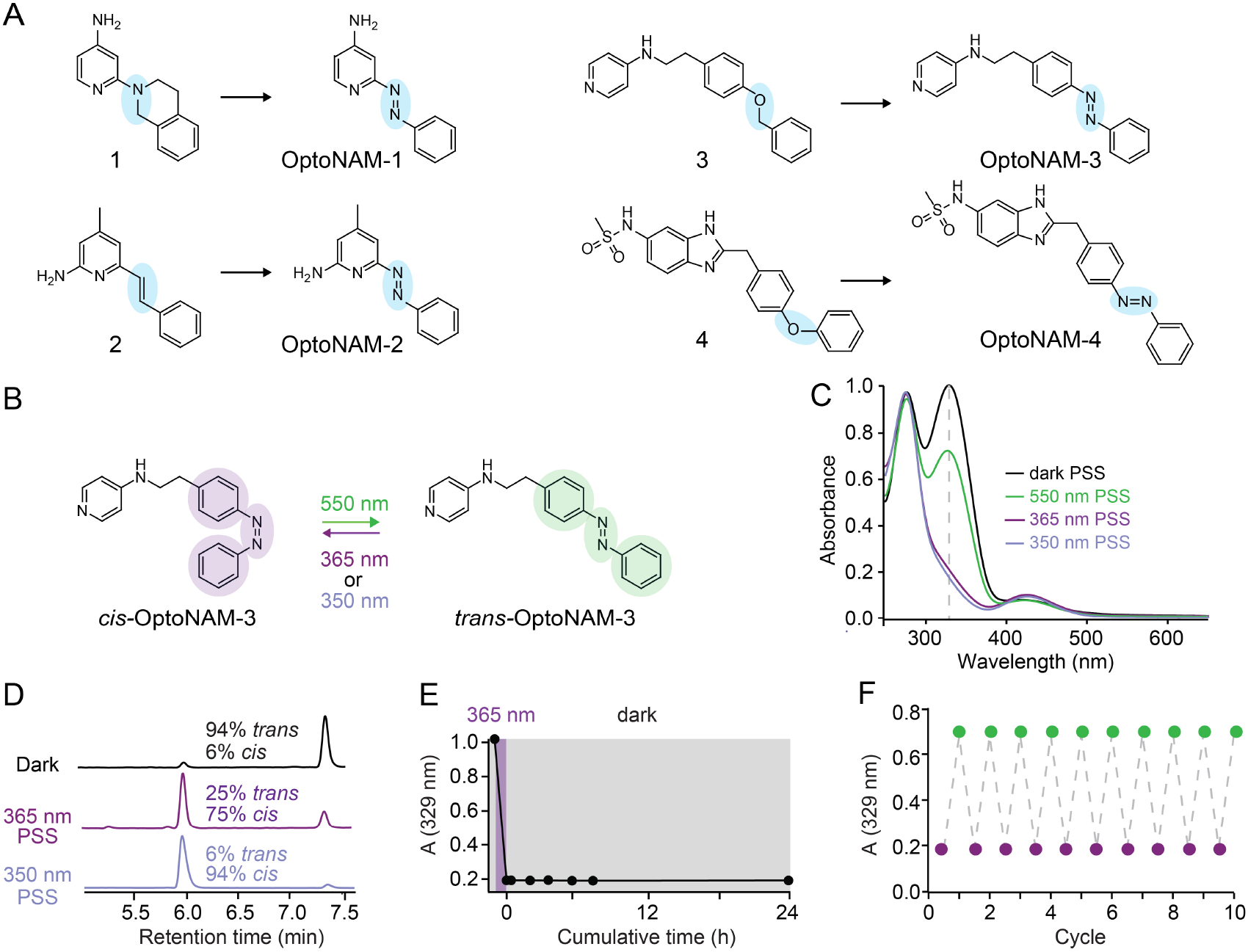
Design and photochemical properties of photoswichable NAMs for GluN1/GluN2B receptors. **(A)** Chemical structures of published GluN2B-selective NAMs (parent compounds 1-4) and their photoswitchable equivalents (OptoNAM-1 to 4) designed by substituting an “azostere” moiety of the parent compound (blue circle) by an azo moiety. Compound **1**, **2**, **3** and **4** correspond to compound **9n** from ref. (54), Compound **14** from ref. (56), Compound **11** from ref (53) and Compound **17a** from ref. (55), respectively. **(B)** OptoNAM-3 can effectively be switched from *trans* to *cis* configuration by UV illumination (365 or 350 nm) and back to *trans* configuration by 550 nm light. **(C)** UV-visible absorption spectra of OptoNAM-3 in the dark, after UV illumination by 350 nm or 365 nm light and after 550 nm illumination of the 365 nm PSS. The dashed line represents the peak absorption wavelength of *trans*-OptoNAM-3 (329 nm). **(D)** HPLC chromatograms of OptoNAM-3 in the dark, and after 365 nm and 350 nm illumination. The photostationary states (PSS) at these different wavelengths were quantified and written next to the peaks corresponding to each isomer. **(E)** OptoNAM-3 365 nm PSS (75% *cis*) is highly photostable in the dark, no change of its absorption spectrum was observed up to 24 h after 365 nm illumination. **(F)** OptoNAM-3 can undergo 10 cycles of UV/green light without degradation. The graph represents the absorbance at 329 nm for each light condition (purple dots for OptoNAM-3 365 nm PSS and green for 550 nm).

**Fig. 2.**
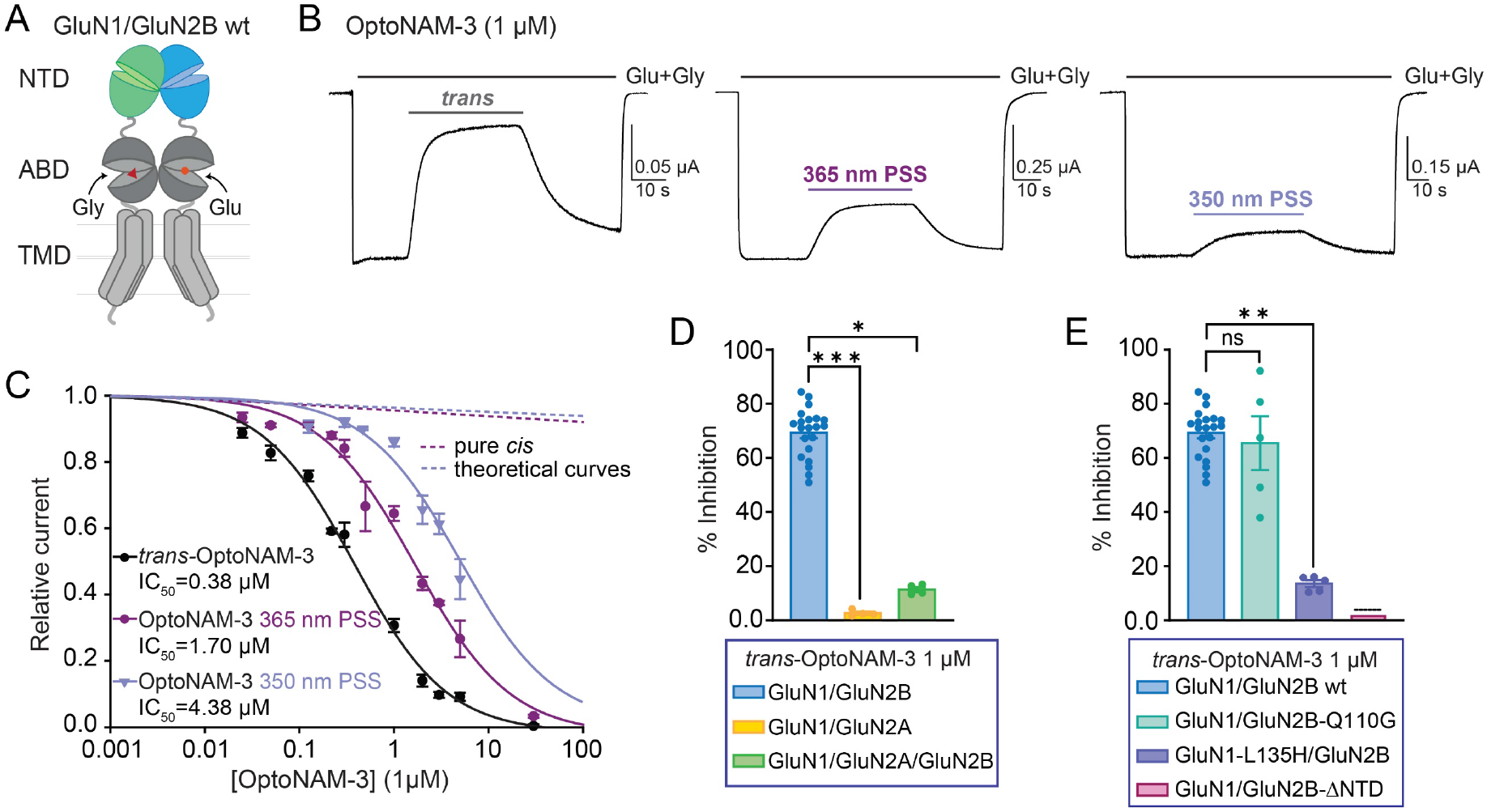
OptoNAM-3 inhibits GluN1/GluN2B selectively and in a photodependent manner. **(A)** Architecture of a dimer of GluN1 and GluN2B subunits with N-terminal domains (NTDs) in green and blue, respectively. ABD, agonist-binding domain; TMD, transmembrane domain. **(B)** Representative current traces from oocytes expressing GluN1/GluN2B receptors following application of agonists glutamate and glycine (Glu + Gly, 100 µM each) and 1 µM OptoNAM-3 either in the dark (left), or pre-illuminated at 365 nm (middle) or at 350 nm (right). **(C)** Dose-response curves of OptoNAM-3 activity on GluN1/GluN2B receptors in the dark (black curve, IC_50_ = 0.38 ± 0.03 µM, n = 5-21) or pre-illuminated with 365 nm (violet curve, IC_50_ = 1.7 ± 0.2 µM, n = 5-17) or 350 nm (lavender curve, IC_50_ = 4.4 ± 0.6 µM, n = 4-13). Theoretical curves of pure *cis-*OptoNAM-3 were calculated either from the 365 nm PSS (25% *trans* / 75% *cis*, violet dotted line) or from the 350 nm PSS (6% *trans* / 94% *cis*, lavender dotted line). **(D)** Comparative inhibitions by 1 µM *trans-*OptoNAM-3 of GluN1/GluN2B and GluN1/GluN2A diheteromers, and GluN1/GluN2A/GluN2B triheteromers (inhibition: 69 ± 2%, n = 21 for GluN1/GluN2B; 2.7 ± 0.5%, n = 4 for GluN1/GluN2A and 11.4± 0.9%, n = 4 for GluN1/GluN2A/GluN2B). n.s., p > 0.05; *, p < 0.05; ***, p < 0.001; Kruskal-Wallis followed by Dunn’s multiple comparison test. **(E)** Percentage of inhibition by 1 μM *trans-*OptoNAM-3 in the dark of wt GluN1/GluN2B receptors (in blue) or of receptors mutated at key residues involved in EVT-101 binding only (GluN1/GluN2B-Q110G, in aquamarine), in ifenprodil binding only (GluN1-L135H/GluN2B, in violet), or on a mutant disrupting binding at both EVT-101 and ifenprodil binding sites (GluN1/GluN2B-ΔNTD, in pink; dotted lines above the bar means that inhibition was calculated from the dose-response curve in Fig. S3C). n.s., p > 0.05; **, p < 0.01; Kruskal-Wallis followed by Dunn’s multiple comparison test.

Photopharmacology, an approach based on the use of photosensitive ligands, allows such high temporal resolution of action. The most widely adopted method for optical modulation of ion channel activity involves caged compounds, whose activity is inhibited by a photolabile moiety (cage) but this strategy is limited by its irreversibility (40, 41). An alternative approach employs photoconvertible ligands that can alternate between an active and an inactive configuration after exposure to different light wavelengths (40–42). This photoswitchable property is conferred to the molecule by the presence in its structure of a photo-isomerisable unit such as an azobenzene, which can reversibly alternate between an extended *trans* form and a twisted *cis* configuration using two different wavelengths, usually UV and blue-green light (40, 43). In a previous paper, we had developed a caged and a photoswitchable ifenprodil derivative (44). However, while caged ifenprodil allowed fast GluN2B-NMDAR inhibition, the kinetics of NMDAR recovery from inhibition were still limited by the slow dissociation rate of ifenprodil. Additionally, the strategy we utilized to obtain a photoswitchable ifenprodil– attachment of an azobenzene to the ifenprodil molecule – strongly decreased its inhibitory activity, probably because ifenprodil binding-site is too small to accommodate the supplementary azobenzene moiety (44).

In this paper, we took advantage of the chemical diversity of GluN2B-selective antagonists (5, 13–15, 24, 45) and designed photoswitchable NAMs by incorporating the azobenzene moiety within the chemical scaffold of the molecule (azologization approach, see refs. (46, 47) and below). Among the four photoswitchable NAM candidates, OptoNAM-3 appeared as a potent and selective inhibitor of GluN2B-NMDARs in its *trans* configuration, while its *cis* isomer was inactive. OptoNAM-3 allowed us to reversibly control GluN2B-NMDAR activity in real time with fast (in the second range) temporal resolution. Surprisingly, binding of OptoNAM-3 to GluN2B-NMDARs induced a red-shift of its action spectrum, allowing us to use visible (blue and green) light to turn off and on OptoNAM-3 activity. Using molecular dynamics and density-function theory (DFT), we show that this red-shift is due to geometric constraints imposed by OptoNAM-3 binding-site on a specific rotamer of the compound. We finally show that OptoNAM-3 also acts as a red-shifted photomodulator *in vivo*, allowing us to reversibly modulate the locomotion behavior of Xenopus tadpoles. This highlights the strong potential of OptoNAM-3 for fast and reversible control of GluN2B-NMDAR activity *in vivo*.

## Results

### Design of a photoswitchable GluN1/GluN2B-selective NMDAR antagonist

There are two main strategies for converting conventional ligands into photoswitchable ligands (46, 47): (i) attach a ’photoswitchable’ moiety (in our case, an azobenzene moiety) to a well-known ligand (azo-extension approach). As this approach can potentially be applied to any ligand, it is currently the most widely used strategy (40–42, 46, 48). (ii) Integrate the azobenzene moiety into the molecular scaffold of the ligand when the latter allows it, a process called “azologisation” (46, 47). This strategy cannot be applied to all ligands and is therefore less employed, although a strong increase in its development has been observed in recent years (46, 47, 49–52). Where appropriate, this technique allows light sensitivity to be conferred on a ligand with minimal chemical modification. We have previously shown that the strategy of azo-extension was not the most appropriate to confer photosensitivity to ifenprodil derivatives because the binding-site was likely too buried to accommodate both the ligand and the azobenzene moiety (23, 24, 44) (see above). We therefore decided to use the azologization strategy. We selected a series of four NAMs selective for GluN2B-NMDARs (13, 53–56) possessing chemical motifs that can be changed into an azobenzene with minimal perturbation of the molecule structure (isosteres of azobenzenes or “azosteres”, refs (46, 47)) to design photoswitchable compounds OptoNAM-1, -2, -3 and -4 (Fig. 1A and Table S1). The photochemical and biological characterizations of OptoNAM-1, -2 and -4 are described in Supplementary Text 1, Fig. S1, S2 and Table S1. In brief, OptoNAM-1 and -2 displayed photodependent activity on GluN1/GluN2B NMDARs but with a >1000-fold shift in IC_50_ compared to their parent compounds (Fig. S1 and Table S1). This is likely due to the loss of protonation of the aminopyridine moiety at physiological pH induced by introduction of the azo moiety (Fig. S2). Protonation was indeed shown to be critical for the activity of this class of compounds onto GluN1/GluN2B NMDARs (Fig. S2D and ref. (54)). OptoNAM-4, on the other hand, retained strong potency for GluN1/GluN2B NMDARs but its activity was not photodependent (Fig. S1 and Table S1). In this paper we focus on OptoNAM-3 (Fig. 1B), which emerged as the best candidate for efficient photocontrol of GluN1/GluN2B NMDARs (see below).

### Photochemical characterization of OptoNAM-3

We first characterized the photochemical properties of OptoNAM-3. We focused on the three key properties allowing use of azobenzenes in biological systems: (i) ideal wavelengths for *trans* to *cis* and *cis* to *trans* conversions; (ii) compound photostability, *i.e.* half-life of the *cis* isomer in the dark; and (iii) fatigability, *i.e.* the number of illumination cycles that the molecule can undergo without degradation. To this aim, we acquired UV-visible absorption spectra of OptoNAM-3 diluted in oocyte recording solution (pH 7.3; see Methods) either in the dark (absence of illumination), or after illumination by light wavelengths ranging from 350 to 580 nm (Fig. 1C and S1M). The absorption spectra of OptoNAM-3 in solution were characteristic of azobenzenes (43). In the dark (black curves in Fig. 1C), the spectrum was characteristic of a *trans* isomer (43). Application of UV light at a wavelength close to the main absorption peak of the *trans* form (365 or 350 nm) gave a completely different spectrum (violet curves in Fig. 1C), characteristic of azobenzenes in their *cis* configuration (43). HPLC analysis of an OptoNAM-3 solution identified photostationary states (PSS) containing 94% *trans* and 6% *cis* in the dark, 75% *cis* and 25% residual *trans* after illumination at 365 nm (365 nm PSS, violet curve in Fig. 1D), and 94% *cis* and 6% residual *trans* after illumination at 350 nm (350 nm PSS, lavender curve in Fig. 1D). This shows that illumination with UV light can convert most of OptoNAM-3 into its *cis* configuration, 350 nm being the most efficient wavelength to induce *trans*-to-*cis* transition. Irradiation of OptoNAM-3 365 nm PSS with wavelengths from 435 to 550 nm gave equivalent *cis*-to-*trans* conversions with 435 and 550 nm yielding slightly stronger conversions (∼70% *trans* after illumination at 435 and 550 nm measured at λ*_trans_* = 329 nm; Fig. 1C and Fig. S1N). 550 nm was chosen as the optimal wavelength since green light is less harmful for cells compared to lights of shorter wavelengths. We then tested the photostability of the *cis* form in the dark (in the absence of illumination). The absorbance spectrum of the 365 nm PSS kept in the dark did not evolve after 24h (Fig. 1E) indicating a very strong photostability of the *cis* isomer in aqueous solution in the dark. This very high stability is interesting because it avoids the prolonged use of UV to keep OptoNAMs in *cis*. OptoNAM-3 could finally endure many illumination cycles without degradation. Indeed, UV-visible spectra acquired after alternating illumination of OptoNAM-3 with UV light and green light during ten cycles were perfectly superimposable for each wavelength (Fig. 1F).

### OptoNAM-3 is a potent NAM of GluN1/GluN2B NMDARs with a photodependent activity

The activity of OptoNAM-3 was then monitored by electrophysiology on Xenopus oocytes (see Methods). To assess the light-dependent effect, we tested the activities of the dark PSS (mostly *trans*) and the UV PSS (mostly *cis*, see above) separately. For each concentration, we made one solution that was divided into two samples: one in which the compound was kept in the dark for the duration of the experiment (dark PSS state), and one that was pre-illuminated with UV light (350 or 365 nm: UV PSS) and then kept out of the light for the duration of the experiment to avoid photoisomerization of the compounds by ambient light (see Methods). Due to their high photostability, the *cis*-OptoNAM-3 did not show any relaxation to the *trans* state during the several hours of experimentation. We generated dose-response curves of OptoNAM-3 dark and UV PSS on wild-type (wt) GluN1/GluN2B NMDA receptors (Fig. 2A) in the presence of saturating concentrations of agonists (glutamate and glycine). OptoNAM-3 had an IC_50_ of 380 nM (Fig. 2B, C) in the dark, which is in the same range of activity as its parent compound **3** (Ki = 93 nM, ref. (53); Table S1). In addition, OptoNAM-3 displayed a significant photo-dependent activity, since its IC_50_ for GluN1/GluN2B NMDARs increased by 4.5 and 11.5-fold compared to the dark condition when the solution was pre-illuminated with 365 and 350 nm light, respectively (IC_50_ = 1.7 µM and 4.38 µM at 365 and 350 nm, respectively; Fig. 2B, C and Table S1). To gain further insights into the photo-dependence of OptoNAM-3 activity, we calculated the theoretical dose-response curve of a pure *cis*-OptoNAM-3 population, knowing that solutions pre-illuminated with 365 nm and 350 nm contain respectively 25% and 6% of residual *trans*-OptoNAM-3, and assuming that the dose-response curve in the dark represents the activity of a pure *trans* population (see Fig. 1D, Fig. 2C and Methods). Our calculations show that the *cis* isomer is inactive on GluN1/GluN2B NMDARs and that the residual activity observed after UV illumination entirely results from the activity of the remaining *trans* isomer (Fig. 2C and S3A, B). The activity of OptoNAM-3 is therefore strongly photo-dependent with an active *trans* isomer and an inactive *cis* isomer. The separation of effect between dark and UV conditions is only limited by the photochemical properties (i.e. the proportion of *trans* remaining after UV illumination) of the compound.

NMDARs exist as multiple subtypes in the brain that are formed by the combination of two GluN1 and either two identical (di-heteromers) or different (tri-heteromers) GluN2 (GluN2A-D) subunits (1). We assessed the selectivity of *trans-*OptoNAM-3 (dark PSS) for the other most abundant NMDAR subtypes in the adult forebrain aside from GluN1/GluN2B di-heteromers: GluN1/GluN2A di-heteromers and GluN1/GluN2A/GluN2B tri-heteromers. 1 µM of OptoNAM-3, which induces 69% inhibition of GluN1/GluN2B diheteromeric NMDARs, induced minimal inhibition (2.7%) of GluN1/GluN2A diheteromers and had slightly stronger effect on GluN1/GluN2A/GluN2B triheteromeric NMDARs (11.4% inhibition, Fig. 2D). Like other ifenprodil derivatives (57, 58), OptoNAM-3 is therefore selective for GluN2B-containing NMDARs with a marked preference for GluN1/GluN2B di-heteromers over GluN1/GluN2A/GluN2B tri-heteromers.

We finally investigated the location of OptoNAM-3 binding-site on the receptor. GluN2B-selective antagonists like ifenprodil are known to bind at the interface between GluN1 and GluN2B NTD upper lobes (23, 26). *Trans-*OptoNAM-3 activity is, like ifenprodil, drastically reduced in receptors in which the NTD of GluN2B has been deleted (59, 60) (GluN1/GluN2B-ΔNTD receptors, 100-fold shift in IC_50_ between wt and GluN1/GluN2B-ΔNTD receptors, Fig. 2E and S3C). In addition, *trans*-OptoNAM-3 IC_50_ was increased in the presence of ifenprodil, which is consistent with a competition between the two compounds (Fig. S3D). The binding site for GluN2B-selective antagonists at the GluN1/GluN2B NTD dimer interface contains two partially overlapping pockets that accommodate GluN2B-selective NAMs of distinct chemical scaffolds (24): either compounds with scaffolds related to ifenprodil, or compounds with scaffolds related to another GluN2B-selective NAM called EVT-101 (61). By mutating residues selectively disrupting binding of the compounds in one or the other pocket (24), we show that OptoNAM-3 binds the ifenprodil binding pocket and not the EVT-101 pocket (Fig. 2E). We have therefore designed a potent NMDAR NAM, OptoNAM-3, which shares the same binding site and selectivity for GluN2B-containing NMDARs as previous GluN2B-selective antagonists but, in addition, displays a strong photodependence of effect, with the *trans* isomer being the only active form on GluN2B-NMDARs.

### Fast and reversible photomodulation of GluN2B-NMDARs in mammalian cells

Now that we have established the photo-dependence of OptoNAM-3 action, we tested whether this compound could be used to perform real-time modulation of NMDAR activity with light. To answer this question, we turned to mammalian cells, whose transparency allows homogeneous illumination of all membrane-expressed NMDARs. OptoNAM-3 was perfused during application of agonists on HEK cells expressing GluN1/GluN2B receptors. When applied in the dark, 2 μM OptoNAM-3 induced 77% inhibition of GluN1/GluN2B currents (Fig. 3A, B). This inhibition was partially abolished by UV (365 nm) illumination (23% remaining inhibition) and partially restored by 550 nm illumination (59% inhibition) (Fig. 3A, B). OptoNAM-3 thus allows real-time reversible inhibition of GluN1/GluN2B activity with light. We furthermore plotted the effect of different concentrations of OptoNAM-3 in the dark and during UV illumination and observed a 20-fold UV-induced shift of OptoNAM-3 IC_50_ compared to the dark condition (Fig. 3C). This shift was greater than when the compound was pre-illuminated in solution and then applied onto oocytes (see above Fig. 2C; 4.5-fold shift in IC_50_ between the dark and the 365 nm conditions). The stronger photodependence of OptoNAM-3 action on HEK cells might either stem from differences between cellular expression systems (Xenopus oocytes vs HEK cells), or from the different irradiation contexts (in solution for experiments in Xenopus oocytes and in a cellular context for HEK cells) (see below)

**Fig. 3.**
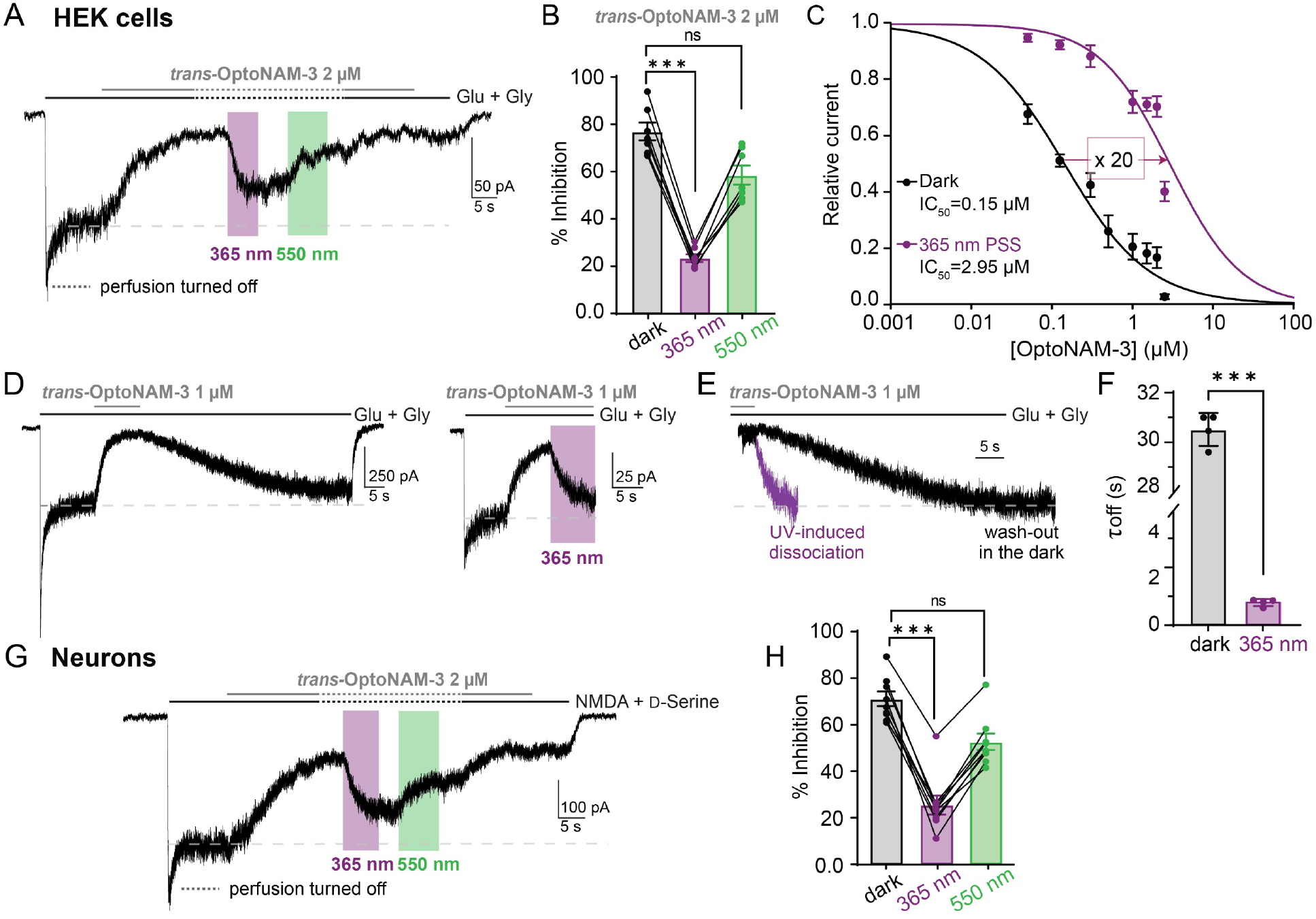
Fast and reversible photomodulation of GluN1/GluN2B receptors in mammalian cells. **(A)** Current trace from a HEK cell expressing GluN1/GluN2B receptors following application of glutamate and glycine (Glu + Gly, 100 µM each) and 2 µM OptoNAM-3 in the dark (mostly in *trans*). Once steady-state inhibition by *trans*-OptoNAM-3 was reached, perfusion was stopped (dashed lines) and the patched cell and the surrounding extracellular medium were illuminated by 365 nm (violet bar) then 550 nm light (green bar). **(B)** OptoNAM-3 inhibition depending on the light conditions. On average, 2 µM OptoNAM-3 inhibits GluN1/GluN2B currents by 77 ± 4% in the dark, 23 ± 1% under UV light and can be restored to 59 ± 4% by 550 nm light. n = 7; n.s., p > 0.05; ***, p < 0.001; Friedman (paired) test followed by Dunn’s multiple comparison test. **(C)** Dose-response curves of OptoNAM-3 on GluN1/GluN2B expressed in HEK cells in the dark (black curve, IC_50_ = 0.15 ± 0.03 µM, n = 4-8) and when the cell is illuminated at 365 nm (violet curve, IC_50_ = 3.0 ± 0.9 µM, n = 4-7). **(D)** Current traces showing the relief of *trans*-OptoNAM-3 inhibition by OptoNAM-3 washout in the dark (left) or by UV light illumination (right). **(E)** Superposition of the inhibition relief traces in the dark (black) and under UV light (violet) showing much faster relief of inhibition by UV illumination than washout of *trans*-OptoNAM-3. **(F)** Summary of the kinetics of inhibition relief in the dark and UV conditions (τ_off_ = 30.5 ± 0.5 s, n = 4 in the dark and τ_off_ = 0.82 ± 0.06 s, n = 4 under UV illumination). ***, p < 0.001; Mann-Whitney test. **(G)** Current trace from a DIV 8 cultured cortical neuron following application of NMDA (300 µM) and D-serine (50 µM), and 2 µM OptoNAM-3 in the dark (mostly in *trans*). Same protocol as in panel (A). **(H)** OptoNAM-3 inhibition depending on the light conditions. On average, 2 µM OptoNAM-3 inhibits NMDA currents by 70 ± 3% in the dark, 25 ± 4% under UV light and can be restored to 52 ± 3.5% by applying a 550 nm light. n = 9, n.s., p > 0.05, ***, p < 0.001; Friedman test followed by Dunn’s multiple comparison test.

The dissociation rates of compounds acting at the ifenprodil site are usually very slow (in the minute range) (34–37). This slow dissociation rate is actually an issue in native tissues like brain slices, in which the effects of such compounds become irreversible. The dissociation rate of *trans*-OptoNAM-3, monitored by the time constant of current recovery from inhibition after OptoNAM-3 washout in the dark, was similarly slow (τ_off_ = 30.5 ± 0.5 s, Fig 3D-F). Interestingly, relief of OptoNAM-3 inhibition by UV light in HEK cells was much faster with a time constant in the sub-second time-range (τ_off_ = 0.82 ± 0.06 s, Fig. 3D-F).

We repeated these experiments on cultured cortical neurons at Days In Vitro [DIV] 6-8, a stage at which GluN2B-containing NMDARs form the major population of neuronal NMDARs (62, 63). We obtained similar results: 70% inhibition of NMDA-induced current by 2 µM OptoNAM-3 in the dark, which was decreased to 25% under UV light illumination and restored to 52% by 550 nm light (Fig. 3G, H). The lower inhibitory effect of OptoNAM-3 on neurons compared to HEK cells is most likely due to the mixture of NMDAR subtypes expressed in neurons (62, 63). With its strong photodependence of action and its fast kinetics of photomodulation, OptoNAM-3 should thus allow fast and reversible inhibition of native GluN1/GluN2B receptors, something that was not possible with regular GluN2B-selective NAMs.

### OptoNAM-3 acts as an *in-situ* red-shifted photodependent antagonist

Given the slow dissociation rate of OptoNAM-3 in its *trans* configuration (dark condition), we hypothesized that the fast relief of inhibition observed upon UV-light illumination resulted from *trans*-to-*cis* interconversion of OptoNAM-3 inside its binding site. The UV-induced relief of inhibition might then reflect either dissociation of *cis*-OptoNAM-3 from the binding site, or the isomerization rate from the active *trans* to the inactive *cis* with the *cis* remaining in the binding site (silent modulator). To further investigate the mechanisms by which this compound exerts its photodependent biological activity, we studied the spectral dependence of OptoNAM-3 photoisomerization in solution (referred to “free OptoNAM-3” below) and in a cellular context (referred to “bound OptoNAM-3” below). Azobenzenes indeed exhibit strong electronic absorption of their conjugated pi system and their absorption spectra can be altered when they aggregate, are complexed, or simply dwell in a different solvent (64, 65). We would thus expect the photochemical properties of bound OptoNAM-3, which is confined in its binding site and exerts multiple non-bonding interactions with it, to differ from the ones of free OptoNAM-3.

To compare the photochemical properties of free and bound OptoNAM-3, we measured the degree of OptoNAM-3 photoisomerization when illumination was performed either on cultured neurons pre-equilibrated with OptoNAM-3 (bound OptoNAM-3, Fig. 4A-C and F-H) or in solution (free OptoNAM-3, previously done by UV-visible spectral analyses, Fig. 1C and Fig. 4D and I). We first analyzed OptoNAM-3 *trans*-to-*cis* isomerization (Fig. 4A-E). To this aim, OptoNAM-3 in the dark was irradiated with light of various wavelengths. The degree of photoisomerization of bound OptoNAM-3 was calculated from the percentage of inhibition induced by OptoNAM-3 under the different wavelength conditions (Fig. 4A-C, E and see Methods). The degree of free OptoNAM-3 photoisomerization was calculated by UV-visible spectroscopy, by measuring the absorbance of the irradiated solution at *trans*-OptoNAM-3 peak absorption wavelength (Fig. 4D, E and see Methods). We observed that wavelengths up to 460 nm allowed efficient recovery from inhibition by *trans*-OptoNAM-3 on cortical neurons (52% inhibition, corresponding to ∼68% of *cis* for the 460 nm PSS; Fig. 4A, C, E), whereas 435 and 460 nm illuminations allowed only partial *trans*-to-*cis* isomerization in solution (∼27% of *cis* after 460 nm illumination, Fig 4D, E). After calculation of the amount of *trans*-to-*cis* photoisomerization of bound and free OptoNAM-3 for all the wavelengths tested (Fig. 4E and see Methods), we observed a red-shift of the action spectrum of OptoNAM-3 in the binding site compared to in solution: wavelengths up to 460 nm allowed a good *trans*-to-*cis* photoconversion of bound OptoNAM-3, while for free OptoNAM-3, *trans*-to-*cis* conversion was unfavored for wavelengths superior to 380 nm. Interestingly, we also calculated a better *trans*-to-*cis* isomerization of bound OptoNAM-3 by 365 nm light (∼90% cis for bound OptoNAM-3 365 nm PSS vs 75% cis for free OptoNAM-3 365 nm PSS, Fig. 4E), which is consistent with the better separation of OptoNAM-3 activity between the dark and 365 nm conditions when the compound was directly irradiated on the cell (Fig. 3C) than when it was pre-irradiated in solution (Fig. 2C). On the other hand, wavelengths of 550 and 580 nm did not induce visible *trans*-to-*cis* isomerization in the binding site, while they induced a significant photoconversion in solution (24 and 22% of *cis* after illumination respectively) (Fig. 4B, E).

**Fig. 4.**
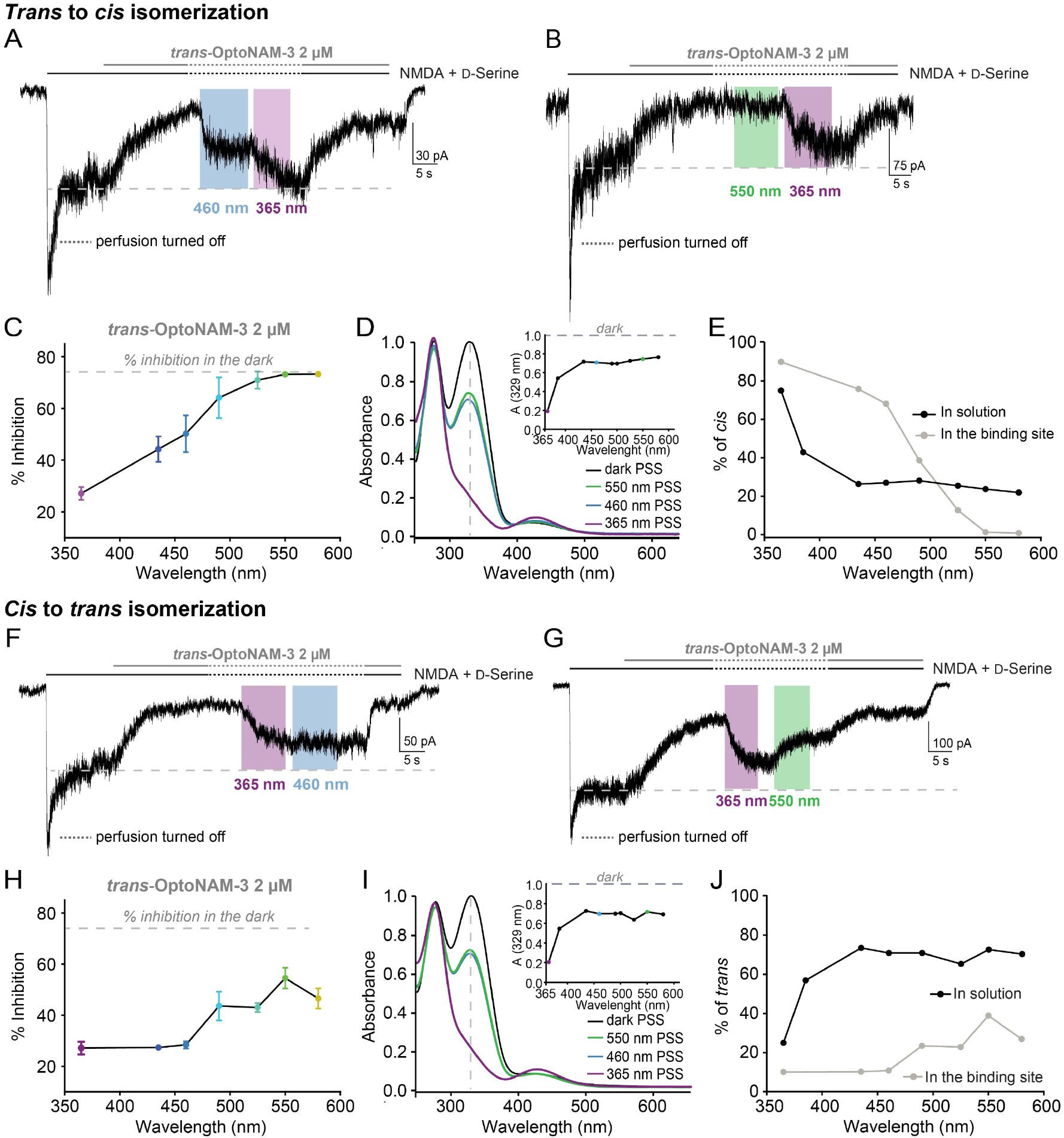
Binding of OptoNAM-3 onto GluN1/GluN2B NMDARs changes its photochemical properties. (A-E) Spectral properties of *trans* to *cis* isomerization. **(A, B)** Current traces from DIV 6-8 cultured cortical neurons following application of NMDA (300 µM) and D-serine (50 µM), as well as 2 µM OptoNAM-3. As described in Fig. 3, once steady-state inhibition by *trans*-OptoNAM-3 was reached, the patched cell and the surrounding extracellular medium were illuminated by various wavelengths: in our example 460 nm (blue bar) in (A) and 550 nm (green bar) in (B) followed by 365 nm (violet bar) as a control for the lowest amount of inhibition. **(C)** Percentage of inhibition by 2 µM OptoNAM-3 upon shining different wavelengths on the cell. 365 nm was the most efficient wavelength allowing reduction of inhibition to 27 ± 2.5%. n = 23 for 365 nm; n=4 for 435 nm; n=5 for 490 and 550 nm; n=6 for 460 and 525 nm. Data points from 435 and 580 nm represented in this figure were scaled between 365 nm and dark condition for better accuracy (see Method for calculation protocol). **(D)** UV-visible spectra of a solution of OptoNAM-3 in the dark (black curve) and after illumination with 365, 460 or 550 nm (violet, blue and green curves, respectively). Inset, absorbance at the peak absorbance wavelength of the dark state (329 nm) of OptoNAM-3 following illumination at various wavelengths of the dark PSS. Note that in solution, the 460 nm and 550 nm PSS are similar. **(E)** Proportion of *cis* isomer in OptoNAM-3 PSS for different illumination wavelengths upon irradiation in solution (calculated from the UV-visible spectra as in Panel D, black points) or in a cellular context (calculated from the percentage of OptoNAM-3 inhibition under various wavelengths as shown in panels (A-C), grey points, see Methods for the calculation protocol). This graph reveals a red shifting of OptoNAM-3 spectral properties in a cellular context. **(F-J) Spectral properties of *cis* to *trans* isomerization. (F, G)** Current traces from cortical neurons following application of NMDA (300 µM) and D-serine (50 µM), as well as 2 µM OptoNAM-3. Once steady-state inhibition by *trans*-OptoNAM-3 was reached (dark), the patched cell and the surrounding extracellular medium were illuminated by 365 nm light to convert OptoNAM-3 to a mostly *cis* configuration. Then various wavelengths were applied to convert OptoNAM-3 back to *trans*: in our examples 460 nm (blue bar) in (F) and 550 nm (green bar) in (G). **(H)** Percentage of inhibition by 2 µM OptoNAM-3 upon shining different wavelengths post 365 nm illumination of the cell. 550 nm is the most efficient wavelength allowing recovery of inhibition to 52 ± 3.5%. n = 23 for 365 nm; n=4 for 435, 460, 490 and 525 nm; n=9 for 550 nm and n=5 for 580 nm. Data points from 435 and 580 nm represented in this figure were scaled between 365 nm and dark conditions (see Method for calculation protocol). **(I)** UV-visible spectra of a solution of OptoNAM-3 in the dark (black curve), after 365 nm illumination (violet curve) and subsequent illumination with 460 (blue curve) or 550 nm (green curve). Inset, absorbance at the peak absorbance wavelength of the dark state (329 nm) of OptoNAM-3 following illumination at various wavelengths of the 365 nm PSS. **(J)** Proportion of *trans* isomer in OptoNAM-3 PSS obtained upon irradiation with various wavelengths of the OptoNAM-3 365 nm PSS either in solution or in a cellular context (same calculation methods as in Panel E). This graph reveals a red-shifted and less efficient *cis*-to-*trans* photoconversion in a cellular context (OptoNAM-3 likely in its binding site) than in solution.

We then analyzed the spectral properties of the reverse reaction: OptoNAM-3 *cis*-to-*trans* isomerization. To this aim, OptoNAM-3 365 nm PSS was irradiated with light of various wavelengths to determine the degree of return to the *trans* state. Similarly to the *trans*-to-*cis* conversion, we observed altered spectral properties of the compound when in solution or bound to the receptor. Indeed, wavelengths like 435 and 460 nm did not allow *cis*-to-*trans* isomerization of bound *cis*-OptoNAM-3, as evidenced by the similar degrees of OptoNAM-3 inhibition at 365, 435 and 460 nm (27% inhibition, corresponding to 10% *trans* isomer; Fig. 4F, H, J). On the contrary, these wavelengths allowed significant return to the *trans* state in solution (74 and 71% of trans isomer, respectively; Fig. 4I, J). In addition, 550 and 580 nm did not allow as good as a return to the *trans* state for bound OptoNAM-3 than for free OptoNAM-3 (Fig. 4G, H, J).

When photoswitched in a cellular context, OptoNAM-3 therefore displays red-shifted properties that likely stem from the compound interaction with the NMDAR protein environment. This feature is interesting, since it means that wavelengths in the visible range like 435 or 460 nm may be used *in vivo* as biocompatible wavelengths to convert the *trans* isomer to *cis* and reduce the harmfulness of high energy wavelength such as 365 nm (see below).

### Origin of the red-shifted properties of bound OptoNAM-3

We wondered about the origin of this drastic shift in OptoNAM-3 spectral properties when the compound is bound to its binding-site. We first measured OptoNAM-3 UV-visible spectra and its photoconversion properties in various solvents in a hope to mimic a more hydrophobic environment within the protein (compared to water). We observed more efficient *trans*-to-*cis* isomerization by 365 nm light in DMSO compared to aqueous solution (365 nm PSS: 90% *cis* in DMSO vs 75% *cis* in Ringer; Fig. 1D and Fig. S4), which might account for the larger separation between the dark and 365 nm dose-response curves for bound-OptoNAM-3 (Fig. 3C) than for free OptoNAM-3 (Fig. 2C). However, none of the solvents tested recapitulated the red-shift in *trans*-to-*cis* and *cis*-to-*trans* wavelength-dependence observed for bound OptoNAM-3, as assessed by a similar proportion of *trans* isomer under 460 and 550 nm light for all tested solvents (Fig. S5).

We therefore turned to molecular dynamics and density function theory (DFT) simulations to directly predict the spectral properties of OptoNAM-3 bound to its binding-site. *Trans*-OptoNAM-3 was docked into the ifenprodil binding pocket using the ifenprodil-bound, GluN1/GluN2B NTD dimer crystal structure 5EWJ (24) (see Methods). After docking, we first performed three simulations of 1 µs with no constraints. They all started from the same structure, but were assigned different initial random velocities during the equilibration procedure (see Methods). We followed the evolution of two dihedral angles during these simulations: one that describes the orientation of the NH from aniline (Fig. 5A), and one that describes the orientation of one of the azo group relative to the central azobenzene phenyl (C-C-N=N dihedral angle; orientation of azo; Fig. 5A). During these simulations, the C-N=N-C angles from the azo group stayed at 180° i.e. OptoNAM-3 did not convert from *trans* to *cis*.

**Fig. 5.**
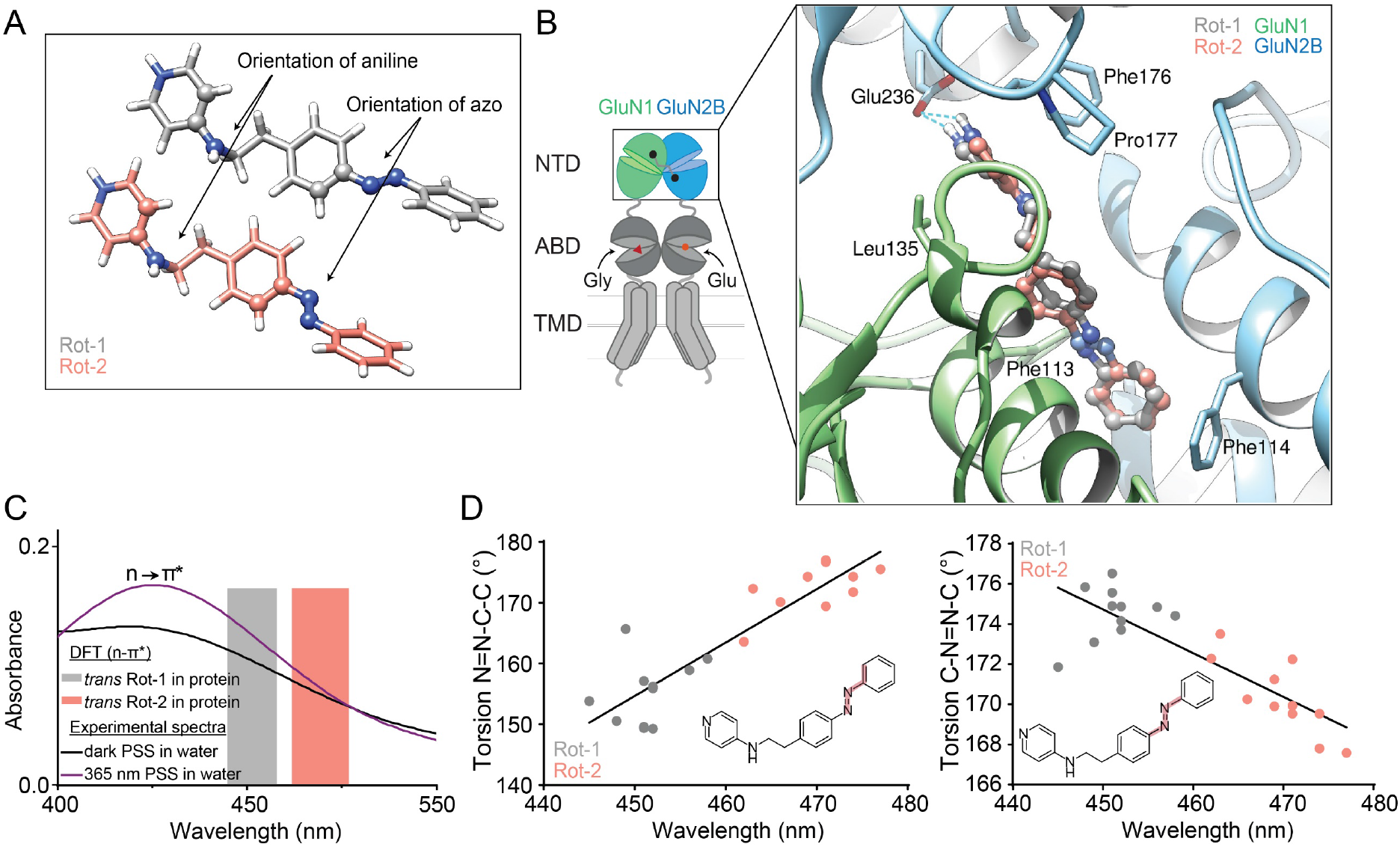
Molecular simulations of OptoNAM-3 in solution and in its binding site. **(A)** 3D representation of *trans*-OptoNAM-3 Rot1 (grey) and Rot2 (salmon) and definition of the two dihedral angles (displayed as balls and sticks) that were monitored during molecular dynamics simulations. **(B)** Docking poses of *trans*-OptoNAM-3 Rot1 (grey) and Rot 2 (salmon) in their binding site: like ifenprodil, at the interface between GluN1 (green) and GluN2B (blue) NTD upper lobes. Residues important for ifenprodil binding (23, 24, 26) are highlighted as sticks. Note that these residues also contact OptoNAM-3 showing a similar binding mode between the two compounds (see also Fig. S10A). **(C)** Superposition of the experimental n→π* band of the UV-Vis spectra of OptoNAM-3 dark PSS (dark line) and 365 nm PSS (violet line), to the theoretical n→π* transitions of OptoNAM-3 *trans* Rot-1 and -2 predicted by DFT calculations (grey and salmon bars representing respectively the range of predicted wavelengths for *trans-*OptoNAM-3 Rot 1 and -2 across the different snapshots of the dynamic, see Table S3). **(D)** Relationships between the N=N-C-C and C-N=N-C torsion angles (as highlighted in pale red in the inset chemical structures) of OptoNAM-3 in the 11 snapshots selected for DFT calculations and their predicted n→π* absorption wavelengths, for Rot-1 (in grey) and Rot-2 (in salmon). This linear correlation shows that a higher torsion angle (lower twist) of the azobenzene leads to a red-shift of the absorption peak.

We observed however that both the aniline and C-C-N=N angles could interconvert between 0° and 180° (Fig. S6A). Thus, the ligand is flexible in the active site with different possible conformations, and in particular different conformations of the terminal phenyl-azo moiety. We furthermore observed that the protein was also quite flexible. Indeed, the distance between GluN1 and GluN2B NTD lower lobes significantly increased during the simulation (35 Å between the Cα of GluN1-K179 and GluN2B-K185 at the end of the simulations vs 25 Å in the 5EWJ crystal structure). This can easily be explained by the fact that we are simulating a dimer of isolated GluN1 and GluN2B NTDs, whose relative orientation is usually constrained by the rest of the protein in the full-length receptor (29). We thus decided to perform new simulations where the protein heavy atoms were restrained close to the crystallographic position by a soft harmonic potential of 100 kJ/mol/nm^2^. One simulation started from the same initial conformation of OptoNAM-3 as before (C-C-N=N angle of 141°, called Rot1; Fig. 5A), whereas in the other we manually rotated the C-C-N=N dihedral angle by 180° (going from 141° to -39°, Rot2 rotamer; Fig. 5A) to reflect the OptoNAM-3 conformational mobility observed in the first round of simulations. As expected, the restraints rigidified the protein, and the distance between GluN1 and GluN2B lower lobes remained around 25 Å. In both simulations, the angle that describes the aniline orientation stayed around 0°, whereas the C-C-N=N angle was respectively of 172° and -17° in average for Rot1 and Rot2, respectively (Fig. 5A, B and Fig. S6B). No transition occurred between the two basins.

To study the behavior of free OptoNAM-3, we performed a 4 µs-long MD simulation of the ligand alone in solution. We observed transitions of both the aniline and the C-C-N=N angle all along the trajectory (Figure S6C). Compared to the simulations performed in the protein, we observed much more transitions between each conformation in solution, which means that the free energy barrier to go from one conformation to the other is much smaller in solution than when bound to the protein.

We extracted 11 snapshots at the end of the simulations of bound- and free-OptoNAM-3 to compute their absorption properties using quantum mechanics (QM) and quantum mechanics/molecular mechanics (QM/MM) calculations (see Methods). We first modeled the absorption properties in water with an implicit scheme where the solvent is represented by a continuum and not by explicit solvent molecules. The computed vertical energies and oscillator strength (f) of OptoNAM-3 are listed in Table S2 using the B2PLYP functional. They are in good agreement with our experimental measurements (Fig. 1C) for the 2 transitions in the visible spectra with a mean deviation of 0.095 eV for the π → π* transition (Fig. S7A and Table S2), validating our computational procedure. We then investigated the absorption properties of the two rotamers of bound OptoNAM-3 (Fig. 5A, B). The computed vertical excitation and oscillator strength (f) of the 11 snapshots are listed in Table S3 for each rotamer. On average the n → π* excitation energy of Rot1 is similar to the one of free-OptoNAM-3 but the n → π* excitation energy of Rot2 is 0.11 eV (20 nm) red-shifted compared to Rot1 (Tables S2 and S3 and Fig. 5C). The second major observation is the difference of the computed oscillator strength (f) of the n → π* between the two rotamers, which is in average 10-fold higher for Rot2 than for Rot1 (Table S3). This dimensionless number can be defined as the probability of absorption in a defined transition. This suggests that the probability of *trans*-*cis* photoconversion can be 10-fold higher for Rot2 than for Rot1.

To investigate the molecular origin of the red-shift in Rot2 absorption, we plotted the relationships between the computed wavelengths of the different snapshots and three different torsion angles of OptoNAM-3: the previously defined C-C-N=N angle; torsion of the azo bond (C-N=N-C); and torsion of the distal azobenzene phenyl relative to the azo bond (N=N-C-C) (Fig. 5D and Fig. S7B-D). We observed a strong correlation between the N=N-C-C torsion angle and the n → π* absorption energy (Fig. 5D). The distal part of the azobenzene moiety was more planar (i.e. closer to 180°) in Rot2, which corresponded to lower excitation energies (higher absorption wavelengths) (salmon dots in Fig. 5D). On the other hand, Rot1 adopted more twisted conformations on this side of the molecule (grey dots in Fig. 5D and S7C), which were closer to the conformations of free OptoNAM-3 in water (blue dots in Fig. S7C). Rot2 isomers furthermore displayed slightly more twisted azo bonds (C-N=N-C angle), which also correlated with a higher absorption wavelength (Fig. 5D and S7B). There was no correlation between the C-C-N=N angle and bound OptoNAM-3 absorption properties (Fig. S7D). It therefore seems that the red-shift of Rot2 n → π* transition in its binding-site is due to geometrical constraints imposed by the protein on the ligand, especially on the extremity of the azobenzene moiety. To confirm this hypothesis, we dissected the electrostatic and geometric contributions of the protein environment to Rot1 and Rot2 transition energies. For both Rot1 and Rot2, re-computing the vertical transition energies in absence of the protein (but keeping the conformations intact) did not significantly affect the compound transition energies (Table S4, lines 1 and 2). On the other hand, removal of the protein and re-optimization of the compound geometries in vacuum induced a 11 nm red-shift of Rot1 transition and a slight, 5 nm blue shift of Rot2 transition, so that the two isomers have similar n → π* transition energies (Table S4, line 3). Rot 1 and Rot2 N=N-C-C angles are much closer to 180° in vacuum (175.41° and 178.89° respectively), which is consistent with our previous analysis correlating a higher torsion angle to a higher n → π* transition wavelength.

In conclusion, the red-shift in bound OptoNAM-3 properties observed experimentally is likely due to geometrical constraints imposed on a specific rotamer of OptoNAM-3, Rot2. While Rot1 adopts similar, twisted conformations whether free or bound to NMDARs, the shape of the binding site imposes a “flatter” conformation to Rot2 of bound OptoNAM-3, hence inducing a red-shift in its n → π* transition. Given the much stronger oscillator strength of Rot2 n → π* transition compared to Rot1, we expect Rot2 to dominate the spectral properties of bound OptoNAM-3.

### OptoNAM-3 controls GluN2B-dependent pathological processes and animal behavior in a red-shifted, photo-dependent manner

We first investigated whether OptoNAM-3 could be used in the context of a patho-physiological process related to GluN1/GluN2B receptors. Overactivation of NMDARs induced by tonic exposure to glutamate or NMDA is well-known to trigger neuronal death (11). GluN1/GluN2B receptors are thought to be the major player of this neurotoxicity and GluN1/GluN2B antagonists have proven to be potent neuroprotectants *in vitro* and *in vivo* (5, 39, 66). To test the neuroprotective activity of OptoNAM-3 and its photo-dependence of action, we performed a protocol of excitotoxicity based on exposure to NMDA (100 µM) of cultured cortical neurons at DIV 14 (66), in the presence of OptoNAM-3. The cell culture was either kept in the dark (OptoNAM-3 in *trans,* active isomer) or illuminated by 365 nm light (converting OptoNAM-3 in mostly *cis*, inactive isomer). OptoNAM-3 in the dark, at a concentration of 5 µM, induced, like ifenprodil, a *∼* 50% increase in cell survival (Fig. S8). This increase in survival was slightly but significantly decreased when UV was applied after addition of OptoNAM-3 (*∼*35% survival in the OptoNAM-3 + UV condition, Fig. S8). No photo-dependent effect was observed on neurons treated with ifenprodil. OptoNAM-3 is therefore able to promote cell survival in neuron cultures in a photo-dependent manner.

We finally demonstrated that OptoNAM-3 allows reversible control of animal behavior with light. Xenopus *laevis* tadpoles can be used as a simple Vertebrate model to validate the activity of photoswitchable drugs applied to neurobiology. Indeed, those tadpoles are transparent allowing good light penetration. In addition, the neuronal pathways underlining the aspects of their swimming behavior have been well characterized (67). In particular, NMDA receptors were shown to play a role in their locomotion pattern (68, 69). Moreover, the ifenprodil binding site is conserved in this species (70) and ifenprodil inhibits Xenopus and mammalian GluN1/GluN2B receptors with similar affinity (71). Xenopus laevis tadpole locomotion is therefore a good behavioral model to test the *in vivo* effects of GluN2B-selective antagonists. We thus tested whether OptoNAM-3 could photomodulate the activity of free-swimming tadpoles.

In our behavioral assay, tadpoles were placed in groups of three animals per well of a 12-well plate allowing them to swim freely. Their baseline locomotion was recorded for three minutes in the dark, after which they were incubated for 45 minutes in a solution containing either 0.1% DMSO (control) or 5 µM OptoNAM-3 (refer to Materials and Methods and Fig. S9A for details). For each condition (control and OptoNAM-3), tadpole locomotion was recorded for three minutes in the dark, followed by cycles of one-minute illumination at 365 nm and 550 nm, interspersed with three-minute rest periods to avoid bias due to a post-stimulation refractory period (67, 72) (Fig. S9A). We also conducted the same protocol using 460/550 nm light cycles, as OptoNAM-3 can switch from *trans* to *cis* state when bound to the receptor upon exposure to blue light at 460 nm (Fig. 4A, E). Tadpole travelled distance was calculated from precise position coordinates of tadpole body parts across frames, assessed by automatic position multi-animal tracking using the open-source program DEEPLABCUT (73, 74). Xenopus *laevis* tadpoles present non visual photoreceptors that generate a photomotor response characterized by an increase of activity upon UV light illumination (75). We indeed observed a small increase of tadpole locomotion with UV compared to 550 nm light in the control condition (up to 1.5-fold, Fig. 6A, B and S9B, C, grey points). The photodependent effect of OptoNAM-3 on locomotion was therefore quantified by calculating the travelled distance normalized twice: by the tadpole baseline locomotion (to decrease inter-individual variability linked to locomotion speed) and, for each condition, by the locomotion of the control group (to remove the unspecific effect of UV light). After incubation with OptoNAM-3, tadpoles from the UV/visible and blue/green cycles protocols (Fig. 6A, C and B, D, respectively) exhibited in the dark a 35% and 50% reduction in locomotor activity, respectively, in comparison to the baseline condition (Fig. 6C, D). Their locomotion could be restored to baseline activity with UV light at 365 nm or blue light at 460 nm and inhibited again with 550 nm for two cycles (Fig. 6C, D). Consequently, OptoNAM-3 enables fast and reversible control of tadpole locomotion using light. In addition, the compound red-shifted properties observed *in vitro* were conserved *in vivo*. OptoNAM-3 therefore holds great promise to photocontrol native GluN2B-NMDARs with the high biocompatibility of red-shifted photoswitches (40, 42).

**Fig. 6.**
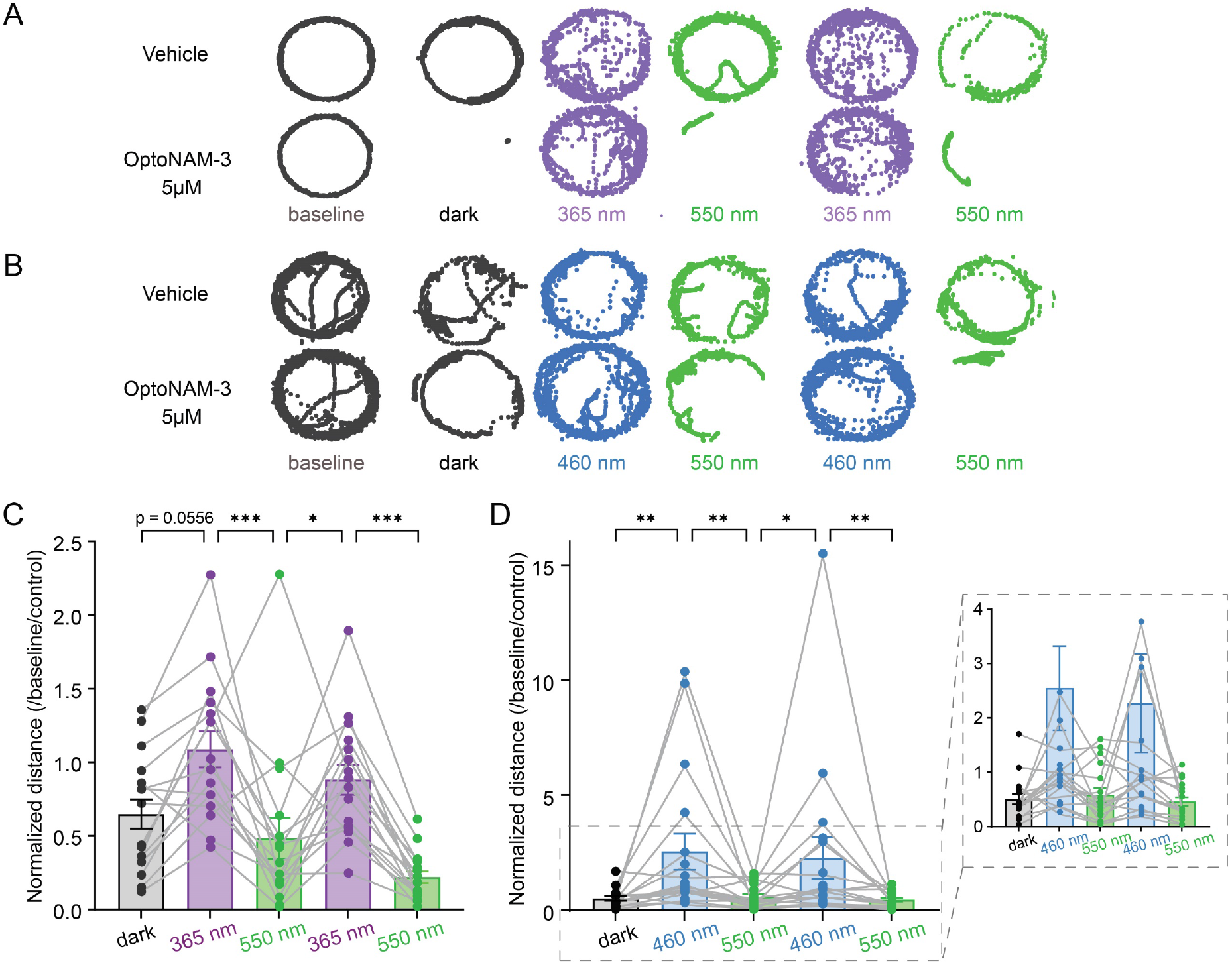
OptoNAM-3 photomodulates Xenopus tadpole locomotion *in vivo*. **(A, B)** Positions (represented by dots) of a tadpole’s stomach in a well taken every 40 ms during 3 minutes of recording for baseline and dark conditions, and 1 minute of recording for either 365 nm and 550 nm illumination cycles (A), or 460 nm and 550 nm illumination cycles (B). Top: tadpole incubated in vehicle (0.1% DMSO, control condition). Bottom: tadpole incubated in 5 µM OptoNAM-3. **(C, D)** Normalized distance traveled by tadpoles (1 point represents the mean traveled distance per well containing 3 tadpoles) exposed to OptoNAM-3 in the dark and during UV/green light cycles (or blue-green light cycles for panel D). n=16 for (C) and n=17 for (D); 48 and 51 tadpoles in total respectively, as the mean of 3 tadpoles per well were calculated to perform a paired statistical test. n.s., p > 0.05; ***, p < 0.001; **, p < 0.01; *, p < 0.05; Friedman test followed by Dunn’s multiple comparison test.

## Discussion

Antagonists selective for GluN2B-NMDARs are routinely used to investigate GluN2B-NMDAR function but their slow rate of action is a strong limiting factor for their use in native tissues. To improve their temporal resolution of action, we have designed a photoswitchable, GluN2B-selective NAM, OptoNAM-3, whose activity could be reversibly turned on and off with light. OptoNAM-3 combined good photochemical properties in solution with efficient *trans*-to-*cis* and *cis*-to-*trans* conversions by UV and blue-green light, respectively; a very strong photostability of its *cis* isomer (>24h); and no signs of bleaching or degradation after 10 cycles of illumination. In the dark (*trans* isomer), OptoNAM-3 combined all the pharmacological properties of GluN2B-selective NAMs: a submicromolar potency (IC_50_ = 0.38 µM) in the same range as ifenprodil and OptoNAM-3 parent compound potencies (53, 60, 76); a strong selectivity for GluN2B-over GluN2A-NMDARs, and the same binding site as ifenprodil derivatives. Inhibition by OptoNAM-3 could be relieved by UV light with a large separation of activity (up to 20-fold) between its dark and UV PSS, which lie in the same range as previously published optopharmacological agents (50, 51, 77). By taking into account the proportion of *cis* and *trans* isomers composing OptoNAM-3 365 and 350 nm PSS, we demonstrated that the remaining inhibition observed after illumination of OptoNAM-3 with UV light was only mediated by the residual *trans* isomer present in the UV PSS. OptoNAM-3 activity is therefore strongly photodependent with one active *trans* isomer and one inactive *cis* isomer. The amplitude of OptoNAM-3 photodependence of effect is thus solely limited by the photochemical properties (i.e. the proportion of *trans* remaining after UV illumination) of the compound.

OptoNAM-3 allowed fast and reversible photomodulation of GluN2B-NMDAR activity in mammalian cells (HEK cells and cultured cortical neurons). In this experimental setting, in which *trans*-OptoNAM-3 was first equilibrated in the cell membrane and inside the GluN1/GluN2B receptor before being irradiated with 365 nm light, we observed a stronger separation of activities between the dark and UV conditions (∼20-fold shift in IC_50_) than when the compound was first UV irradiated and then applied onto the cell (4.5-fold shift in IC_50_, see above). Given our previous results, this suggested that the photochemical properties of OptoNAM-3 in solution and in a cellular context were different. Indeed, by performing a comprehensive comparison of OptoNAM-3 spectral properties (i.e. proportion of *cis* and *trans* isomers under illumination wavelengths from 365 to 580 nm) in solution (free OptoNAM-3) and in a cellular context (bound OptoNAM-3), we observed a strong red shift in the spectral properties of the *trans*-to-*cis* and *cis*-to-*trans* transitions of bound OptoNAM-3 compared to free OptoNAM-3. As a consequence, 460 nm light was almost as efficient as 365 nm light to relieve inhibition by *trans*-OptoNAM-3, while in solution a majority of *trans* isomer was present in the 460 nm PSS. This has important consequences for *in vivo* applications, since use of blue light would be less toxic and have better tissue penetration than UV light (see below). This red-shifting of OptoNAM-3 properties suggests that, when OptoNAM-3 is first equilibrated onto the cell before illumination, photoswitching occurs within the binding site and that the observed red-shifting of OptoNAM-3 photochemical properties stems from the interaction of the molecule with its binding-site.

These changes of OptoNAM-3 spectral properties could not be simply mimicked by changing the polarity of the medium, although we observed a more efficient *trans*-to-*cis* isomerization by 365 nm light in DMSO. Instead, molecular dynamics simulations showed that OptoNAM-3 could bind as two distinct rotamers differing by the orientation of the distal phenylazo moiety (Rot1 and Rot2, Fig. 5A) in the GluN1/GluN2B NTD dimer interface. DFT calculations of the electronic transitions of these two OptoNAM-3 rotamers within their binding-site, revealed that the n→π* transition of the Rot2 isomer was shifted by 20 nm towards higher wavelengths and its absorption coefficient increased by 10-fold. We showed that this red-shift was due to geometrical constraints imposed by the binding site, which stabilized Rot2 azobenzene moiety in a less twisted conformation than in water. Our findings are consistent with previous experiments and simulations showing an increase in n→π* transition energies (and hence a blue-shift in the n→π* absorption bands) as the phenylazo moieties rotate away from planarity (78, 79). This is particularly true for ortho-substituted azobenzenes, in which addition of substituents at the azobenzene ortho positions induces a red-shift in their n→π* transition (78–81), due to electronic repulsions between the nitrogen lone pairs and the ortho-substituents. This red-shift is less important in twisted, ortho-substituted azobenzenes compared to the ones retaining a more planar structure (78, 79). One can thus assume a similar explanation for the origin of the red-shift of bound Rot2 spectral properties. Differences in photochemical properties between free and bound ligands are not well documented in the literature. Such drastic red-shifting of spectral properties has however been previously observed for a photoswitchable AMPA receptor competitive antagonist, ShuBQX-3 (82). The authors attributed this shift to the interaction of ShuBQX-3 azobenzene moiety with an arginine in the inhibitor binding site (82). In their case, free arginine induced a 75 nm red-shift of ShuBX-3 π→π* transition in solution, but the actual impact on the compound spectrum of ShuBQX-3/arginine interaction within the protein was not confirmed by quantum calculations. Given the strong dependence of azobenzene photochemical properties on the molecule geometry and chemical environment, it is highly probable that alterations of the photoswitching properties of photosensitive compounds when bound to their target are more common than currently thought.

While OptoNAM-3 dissociation rate in the dark was slow (τ *∼* 30 s), similarly to other GluN2B-selective NAMs, the reduction of inhibition by UV light was rapid (τ *∼* 1 s). As previously said, the slow dissociation rate of GluN2B-selective NAMs represents a problem when using them in native tissues, in which the effects of the compounds become irreversible. In contrast, the fast light-induced onset and offset of OptoNAM-3 inhibition, in the second time range, should allow dynamic tuning of GluN2B-NMDAR activity *in vivo*. This fast decrease of inhibition is most likely due the *trans*-to-*cis* photo-conversion of OptoNAM-3 inside its binding site. It remains unclear, however, whether the formed *cis*-OptoNAM-3 dissociates from its binding site or remains there as a silent modulator. Photoswitching within the binding site was previously described in mGlu receptors (83, 84), AMPA receptors (82), as well as at the orthosteric glutamate site of NMDARs (85). In the latter study, the authors used this property to induce deactivation of NMDARs with much faster kinetics than after glutamate uncaging (85). Hence, our study confirms that receptor photocontrol using photoswichable ligands is very powerful in that it allows to break the temporal limitations not only of ligand diffusion, but also of ligand association/dissociation.

The 20-fold shift of OptoNAM-3 between dark and UV conditions, its red-shifted properties, as well as the fast kinetics of light-induced GluN2B-NMDAR inhibition and disinhibition observed *in vitro* suggest that our compound should be a powerful tool to dynamically modulate GluN2B-NMDAR activity *in vivo*. Indeed, in the dark OptoNAM-3 inhibited the locomotion of Xenopus tadpoles, a process involving NMDARs (68, 69). Locomotion could be restored to control levels by either application of UV (at 365 nm) or blue light (at 460 nm), and inhibited again by application of visible light (550 nm) for 2 cycles. Our *in vivo* experiments therefore confirm the in-situ, red-shifted properties of OptoNAM-3. Since blue light is less toxic and has better tissue penetration than UV light, OptoNAM-3 is a promising compound to reversibly control GluN2B-NMDAR activity in the mammalian brain.

A large number of soluble, photoswitchable ligands have been currently developed for many different targets including receptors, ion channels and enzymes (see, for instance, refs (40, 42, 48, 83). In iGluRs, the large majority of previously developed photoswitchable compounds are competitive agonists and antagonists (82, 85–88), with a few compounds targeting the ion channel pore (87, 89, 90). However, none of these compounds display subtype selectivity. In this work we have developed the first photoswitchable allosteric modulator of NMDARs, OptoNAM-3, which selectively targets GluN2B-NMDARs. In addition to being more specific than competitive ligands and pore blockers, allosteric modulators preserve the endogenous patterns of receptor activity, since they do not prevent binding of the endogenous agonists (91, 92). This limits the toxicity and desensitization linked to untimed receptor activation/inhibition. OptoNAM-3 combines the high potency and selectivity of classical GluN2B-selective NMDAR antagonists with faster kinetics of action, improved reversibility and red-shifted properties that make it highly biocompatible. This compound should thus allow selective, local and time-controlled inhibition of GluN2B-NMDARs in native systems. It should thus lead to a better understanding of the physio-pathological roles of this class of receptors in the brain, as well as to better tolerated drugs to counteract neurological diseases linked to NMDAR overactivation.

## Materials and Methods

### Chemicals

Salts, D-serine, DTPA (diethylenetriamine-pentaacetic acid), glucose, L-glutamate, glycine, MTT (3-(4,5-dimethylthiazol-2-yl)-2,5-diphenyl tetrazolium bromide), α-cyclodextrin, γ-cyclodextrin, uridine and 5-Fluoro-2’-deoxyuridine were purchased from Sigma-Aldrich (St. Louis, MO, USA); D-APV (D-(-)-2-Amino-5-phosphonopentanoic acid) and N-methyl-D-aspartate from HelloBio (County Meath, ROI) and gentamycin from GIBCO (Invitrogen, Rockville, MD, USA). Ifenprodil is a gift from Sanofi-Synthelabo (Bagneux, France).

OptoNAM-1 (2-(2-phenyldiazen-1-yl) pyridin-4-amine, Mw=198.22), OptoNAM-2 (4-methyl-6-[(E)-2-phenyldiazen-1-yl] pyridine-2-amine, Mw=212.25), OptoNAM-3 (N-{2-[4-(2-phenyldiazen-1-yl) phenyl] ethyl} pyridine-4-amine, Mw= 302.37) and OptoNAM-4 (N-[2-({4-[(E)-2-phenyldiazen-1-yl]phenyl}methyl)-1H-1,3-benzodiazol-6-yl]methanesulfonamide, Mw= 405.47), were obtained from custom synthesis by Enamine Ltd. (Kiev, Ukraine).

Stock solutions of L-glutamate and glycine (100 mM or 1 M each), DTPA (10 mM to 1M at pH 7.3), D-APV (100 mM), were prepared in bi-distilled water. OptoNAMs and ifenprodil stock solutions were prepared by diluting the powder in DMSO to a concentration of 50 mM and 20 mM respectively. They were divided in 0.5 mL aliquots and stored at -20 °C until the day of the experiment.

### Spectroscopic analyses of OptoNAMs

UV-vis absorbance spectra were measured from solutions of 10, 50 or 100 μM free Opto-NAMs diluted in the dark in Ringer (extracellular TEVC recording solution see below) at pH 7.3 in 1 cm long quartz cuvettes, on a NanoPhotometer^®^ NP80 spectrometer (Implen, Germany). Photostationary states of *cis* and *trans* isomers of OptoNAMs in Ringer (pH 7.3, max. 0.1% DMSO) were obtained by continuous illumination with multi-wavelength LEDs (pE-2 and pe-4000, CoolLED, UK (power ∼ 75 mW)); or photochemical reactor RPR-100 Rayonet US, (power ∼ 2 mW) for 350 nm, until no further change in the absorption spectra could be observed. For all irradiation wavelengths tested, 10 min illumination was sufficient to reach steady state.

The *cis-trans* compositions of OptoNAM-3 in the dark and after illumination by 365 and 350 nm were determined by HPLC using the relative integrated areas of the *cis* and *trans* peaks at the isosbestic point (390 nm), considering the sum of integrated area for two peaks is 100%. Analytical HPLC was performed on an Agilent 1200 series equipped with a quaternary pump using a Proto 200 C18 from Higgins Analytical Inc (particles size 3 μm, 100×4.6 mm column). The compounds were eluted with a flow of 1 ml/min using a gradient of acetonitrile (0 to 100% over 10 minutes) in water both solvent containing 0.1% TFA. The detection was performed at 220 nm, 280 nm and 390 nm.

The dark and 365 nm PSS were furthermore characterized using ^1^H-NMR spectroscopy with the OptoNAMs diluted in d^6^-DMSO and the relative abundance of *trans* and *cis* isomers quantified using ^1^H NMR spectra. Indeed, because the aromatic protons of *trans* and *cis* exhibit sufficiently different chemical shifts, their relative abundances at each PSS can be determined by peak integration.

C*is-trans* composition of OptoNAM-3 after illumination at other wavelengths was calculated from their UV-vis spectra using the following formula, considering from the HPLC analysis that the dark PSS is composed of 100% *trans* and the 365 nm state of 25% of *trans*: %*trans* = (A_λ_ – p) / (A_dark_ – p), where p represent the theoretical absorbance of a pure *cis* population p = (A_UV_ - 0.25A_dark_) / 0.75, A_λ_, A_UV_ and A_dark_ the absorbances of the compounds after illumination at a given wavelength, after illumination at 365 nm and in the dark, respectively, measured at the absorption peak of the *trans* state (329 nm).

Photostability of the *cis* state (365 nm PSS) was measured by irradiating the solution with 365 nm light during 10 min, then letting it relax in the dark, inside the spectrophotometer. Spectra were acquired at regular intervals, up to one day, after irradiation.

### Molecular Biology

Rat GluN1-1a (named GluN1 herein), rat GluN2A and mouse GluN2B subunits (ε2) were expressed using pcDNA3-based expression plasmids. The GluN2B-ΔNTD construct is from ref. (59) and GluN2B-N615K from ref. (93). Single mutants GluN2B-Q110G and GluN1-L135H were from ref. (24).

### Oocyte treatment and microinjection

Oocytes from Xenopus *lævis* were used for heterologous expression of recombinant NMDA receptors studied using two-electrode voltage-clamp (TEVC). Female Xenopus laevis were housed and ovary bags harvested according to the European Union guidelines (husbandry authorizations #C75-05-31 and #D75-05-31; project authorizations #05137.02 and Apafis #28867-2020121814485893). Fragments of ovary bags were also purchased from the “Centre de Ressources Biologiques Xenopes” (now TEFOR, Paris Saclay, France) and from the European Xenopus Resource Center (EXRC, Portsmouth, UK). Xenopus *laevis* oocytes were harvested and prepared as previously described in ref. (94).

Expression of recombinant NMDA receptors was obtained by oocyte nuclear co-injection of 37 nL of a mixture of cDNAs (at 30 ng/μL) coding for GluN1*-*1a and various GluN2 subunits (ratio 1:1). The protocol in ref (57) was followed for the expression of GluN1/GluN2A/GluN2B triheteromers. The oocytes were transferred in 96-well plates filled with Barth solution (in mM: 88 NaCl, 1 KCl, 0.33 Ca(NO_3_)_2_, 0.41 CaCl_2_, 0.82 MgSO_4_, 2.4 NaHCO_3_ and 7.5 HEPES, pH adjusted to 7.3 with NaOH), supplemented with gentamicin (50 μg/μL) and 50 μM APV, a selective NMDA receptor antagonist. Plates were then stored at 18 °C for 24h for GluN1/GluN2A expression and 48h to 72h for expression of GluN1/Glu2B, GluN1/GluN2A/GluN2B and mutants.

### Two-electrode voltage clamp and recording solutions

TEVC recordings were performed 1–3 days following injection. TEVC recordings were performed using an Oocyte Clamp amplifier OC-725 (Warner Instruments) and computer-controlled via a 1440A Digidata (Molecular Devices). Currents were sampled at 100 Hz and low-pass filtered at 20 Hz using an 8-pole Bessel filter (900 Series, Frequency Devices Inc). Data were collected with Clampfit 10.3. During the recording, the cells were continuously perfused with external recording Ringer solution at pH 7.3 (in mM: 100 NaCl, 0.3 BaCl_2_, 5 HEPES and 2.5 KCl, pH adjustment to 7.3 by addition of NaOH). The NMDA currents were induced by simultaneous application of L-glutamate and glycine (agonist solution) at saturating concentration (100 μM each), and DTPA (10 μM) to prevent receptor inhibition by ambient zinc (∼20 nM, ref. (95)). A control solution (100 μM glycine and 10μM DTPA in Ringer pH 7.3) was used for washout of drug and test solutions. Unless notified, recordings were performed at a holding potential of -60 mV. All experiments were performed at room temperature.

OptoNAMs were diluted as stock solutions of 50 to 0.5 mM in DMSO. The day of the experiment, they were diluted to the appropriate concentration in the recording solution and kept in the dark during the whole duration of the experiment to avoid photoconversion. Since DMSO itself induces a small inhibition of NMDAR currents, control and agonist solutions were also supplemented with DMSO up to 0.1%. 365 nm PSS solution was obtained by irradiating from the top 25 mL of *trans*-OptoNAM solutions with a 365 nm light LED (pE-2, CoolLED, UK, power ∼ 75 mW) for 10 min in a graduated cylinder covered with aluminum foil. To obtain 350 nm PSS solutions, solutions containing OptoNAM *trans* were put in a quartz cuvette in a photochemical reactor (RPR-100, Rayonet, US, irradiance power ∼ 2 mW) at 5 cm from a 350 nm neon for 15 min. Recordings with 350 nm PSS were performed within 20 min after 350 nm irradiation to avoid photoconversion.

### Whole-cell patch-clamp recordings and photomodulation in HEK cells

HEK293 cells were used for heterologous expression of recombinant NMDA receptors studies using whole-cell patch-clamp. Wild-type NMDARs were expressed in HEK-293 cells (obtained from ECACC, Cat #96121229). HEK cells were cultured in DMEM + glutamax medium supplemented with 10% fetal bovine calf serum and 1% Penicillin/streptomycin (5000 U/ml) and cultured under standard cell culture conditions (95/5% O2/CO2 mixture, 37 °C). Transfections were performed using polyethylenimine (PEI) in a cDNA/PEI ratio of 1:3 (v/v). Cells were co-transfected with a DNA-mixture containing plasmids encoding wild-type GluN1, wild-type GluN2A or GluN2B and eGFP. The total amount of DNA was 1.0 μg per 500 µL of transfected medium containing 12 mm² diameter coverslip and the mass ratio of GluN1:GluN2B:eGFP was 1:2:1 (1:1:1 for GluN2A). 150 µM of D-APV vas added to the culture medium after transfection.

Receptor functionality, OptoNAM-3 photodependence (dark then UV light) of activity as well as association/ dissociation kinetics (with and without UV light) were assessed in patch-clamp recordings of lifted whole cells 24–72 h post transfection (cells were not lifted for the 2 light cycles protocol, i.e. when “perfusion turned off” was written below current traces). Positively transfected cells were visualized by GFP-fluorescence. The extracellular solution contained (in mM): 140 NaCl, 2.8 KCl, 1 CaCl_2_, 10 HEPES, 20 sucrose and 0.01 DTPA (290– 300 mOsm), pH adjusted to 7.3 using NaOH. Patch pipettes had a resistance of 3–6 MΩ (whole-cell) and were filled with a solution containing (in mM): 115 CsF, 10 CsCl, 10 HEPES and 10 BAPTA (280–290 mOsm), pH adjusted to 7.2 using CsOH. Currents were sampled at 10 kHz and low-pass filtered at 2 kHz using an Axopatch 200B amplifier, a 1550B digidata and Clampex 10.6 (Molecular Devices). Agonists (100 μM glutamate and 100 μM glycine) were applied using a multi-barrel solution exchanger (RSC 200; BioLogic). Recordings were performed at a holding potential of −60 mV and at room temperature.

For the c*is* to *trans* isomerization study (Fig. 3A, B): OptoNAM-3 was perfused in the dark onto the patched cell. Once the inhibition reached steady-state, indicating that the compound was in its binding site, perfusion was stopped and 365 nm was applied to the cell. Then, wavelengths from 435 to 580 nm were applied and the inhibition recovery monitored.

Computer-controlled light pulses during electrophysiological recordings were provided from high power sensitive LEDs (CoolLed pe-4000: 4 channels each controlling 4 wavelengths from 365 nm to 770 nm). The LED port was directly coupled via a microscope adaptor to the fluorescence port of an inverted IX73 Olympus microscope. The output beam of the LED entry was directed towards the sample thanks to a mirror (Chroma) and applied to the center of the recording dish through a 10X objective (irradiance ∼ 4 mW/mm² for 365, 385, 435 nm; ∼ 6.5 mW/mm² for 460 nm; ∼ 2 mW/mm² for 490 and 525 nm; ∼ 1.5 mW/mm² for 500 nm and ∼ 4.5 mW/mm² for 550 and 580 nm) (Olympus, 0.30 N.A.).

Light power was measured in the center of the recording dish plane with an optical power meter (1916-C, Newport) equipped with a calibrated UV/D detector and irradiance was obtained upon dividing light power by the illuminated field of the microscope 10X objective (19 mm²).

### Primary cortical neurons cultures

Mice were housed in the IBENS rodent central facility duly accredited by the French Ministry of Agriculture. All experiments were performed in compliance with French and European regulations on care and protection of laboratory animals (EU Directive 2010/63, French Law 2013-118, February 6th, 2013), and were approved by local ethics committees and by the 19 French Ministry of Research and Innovation (authorization numbers #05137.02, APAFIS #28867-2020121814485893 and APAFIS #29476-2021020311595454). Dissociated cultures of cortical neurons were prepared from mouse embryos at E18 as described previously (96). After removing meninges, cortices were placed in ice-cold HBSS solution. Cell dissociation was performed individually for cortices of each embryo. Cortices were incubated in 2.5% trypsin at 37 °C for 7 min, rinsed three times with 37 °C phosphate buffer saline, and finally suspended in plating medium (Neurobasal medium, supplemented with 0.5 mM L-glutamine, 1% B27 supplement and Penicillin/Streptomycine). The neurons were further dissociated by trituration, and cells were plated on poly-D-lysine coated coverslips in 24-well culture dishes at a density of 3*10^5^ cells per well for the excitotoxicity test and 1*10^5^ cells per well for patch clamp experiments. Cultures were incubated at 37 °C in a humidified atmosphere of 95% O2, 5% CO2. Cells were fed by changing ½ medium to fresh Neurobasal medium every 4 days. For the excitotoxicity experiments, after 5 days in vitro, growth of non-neuronal cells was halted by a 24 h exposure to FDX (5 mM uridine and 5 mM 5-Fluoro-2’-deoxyuridine). The cultures were used for experiments after 6 to 14 days in vitro (DIV6-8 for patch clamp experiments and DIV14 for excitotoxicity tests).

### Whole-cell patch-clamp recordings in wild type cultured cortical neurons

Whole-cell patch-clamp recordings on neurons were performed 6 to 8 days after the culturing step (DIV6-8) using the same recording conditions as for HEK cells. Recordings were performed at a holding potential of −60 mV and at room temperature. Currents were elicited by NMDA (300 µM) and D-serine (50 µM) to specifically activate NMDARs.

T*rans* to *cis* isomerization study (Fig. 4A-C): OptoNAM-3 was perfused in the dark onto the patched neuron. Once the inhibition reached steady-state, indicating that the compound was in its binding site, perfusion was stopped and wavelengths from 435 to 580, followed by 365 nm (the wavelength allowing optimal *trans*-*cis* isomerization) were applied. Stopping perfusion ensures that trans-OptoNAM coming from the perfusion tube can compete with the wavelength PSS during the illumination procedure.

C*is* to *trans* isomerization study (Fig. 3G-H and 4F-H): OptoNAM-3 was perfused in the dark onto the patched neuron. Once the inhibition reached steady-state, indicating that the compound was in its binding site, perfusion was stopped and 365 nm was applied to the cell. Then, wavelengths from 435 to 580 nm were applied and the inhibition recovery monitored.

### Neuronal toxicity experiments and assessment of neuronal death

Excitotoxicity tests were performed 14 days after the culturing step as described in ref. (66). Just before the experiments, the neuronal medium was changed and the neurons incubated in the external patch recording solution (see above). Two replicates of 24 well plates containing cultured cortical neurons where exposed to the same 4 conditions, in which the following compounds were added to the medium: (1) glycine (10 µM) alone (control); (2) glycine (10µM) + NMDA (100 µM); (3) glycine (10 µM) + NMDA (100 µM) + 5 µM ifenprodil; and (4) glycine (10µM) + NMDA (100 µM) + 5 µM *trans*-OptoNAM-3 (dark). One of the two plates was exposed to UV light (Jena analytic US, UVP Handheld UV lamp UVGL-58, power ∼ 1 mW) for 2 min. The two plates were then put back in the 37 °C incubator for 10 min. Agonist exposure was terminated by washing out the exposure solution with conditioned Neurobasal medium, prior to returning the dishes to the incubator for 24 h before assessment of neuronal death.

Overall neuronal cell death was determined 24 h after NMDA exposure by the MTT test. Mitochondrial and cytosolic dehydrogenases of living cells reduce the yellow tetrazolium salt (MTT) to produce a purple formazan dye that can be detected spectrophotometrically (97). MTT was dissolved in phosphate-buffered saline (PBS) buffer at 5 mg/ml and filtered through a 0.2 µm membrane to sterilize it and remove a small amount of insoluble residues present in some batches of MTT. A day after the cell exposure to the different conditions, 50 µL of stock MTT was added to all 500 µL medium containing wells, and plates were incubated at 37°C for 4 h. The medium was removed and 200 µL of warm DMSO was added per well and mixed thoroughly to dissolve the dark blue crystals. After 10 min at 37 °C to ensure that all crystals were dissolved, the absorbance of the wells was monitored at 540 nm on a multimode plate reader Infinite 200 PRO R (Tecan, Switzerland).

### Data analysis

Data were collected and analyzed using pClamp 10.5 (Molecular Devices) and fitted using Sigmaplot 11.0 (SSPS). Unless otherwise mentioned, error bars represent the standard deviation of the mean value. Dose-response curves were fitted with the following Hill equation: I_rel_ = 1−a/(1 + (IC_50_/[B])^nH^), where I_rel_=I_antago_/I_control_ is the mean relative current, [B] the drug concentration, (1−a) the maximal inhibition and nH the Hill coefficient. IC_50_, a, and nH were fitted as free parameters. IC_50_ errors represent the error of the fit.

Theoretical “pure *cis”* dose-response curves of OptoNAM-3 (Fig. 2C) were obtained by calculating the theoretical relative current elicited by a pure OptoNAM *cis* population (I_Relcis_) at each concentration (C) data point from the following equation: I_Rel*cis*_=1+(I_RelUVPSS_-I_Rel*trans*_). I_RelUVPSS_ is the relative current obtained from the 365 nm PSS dose-response curve following application of a concentration C of OptoNAM, corresponding to 75% *cis*-OptoNAM and 25% *trans*-OptoNAM (and 94% cis / 6% trans if starting from the 350 nm PSS dose-response curve). I_Rel*trans*_ is the relative current obtained from the *trans*-OptoNAM dose-response curve (dark) following application of a concentration of 0.25xC of *trans*-OptoNAM-3 (or 0.06xC if deduced from the 350 nm PSS dose-response curve).

The theoretical 25% (365 nm PSS *trans*% obtained by HPLC) and 6% of *trans* (350 nm PSS obtained in HPLC) dose-response curves (Fig. S3A, B) were obtained starting from the *trans-*OptoNAM-3 DR curve by considering that the dark state of OptoNAM-3 solution is composed of 100% of the *trans* isomer and that each relative current obtained for one concentration I_RelC_ is equal to the relative current I_Rel%C_ exerted by the application of 25%, 15% or 5% of this concentration.

The kinetics of photoinactivation and OptoNAM-3 dissociation (Fig. 3E, F) were obtained by fitting currents with a single exponential function as follows: Y = A*exp(-t/τ) + C, with A as the initial current amplitude, Tau (τ) the decay time constant, and C the steady-state level.

For the *trans* to *cis* and *cis* to *trans* isomerization study in binding site (Fig 4C and H respectively), we decided to normalize (scale) the inhibition values of OptoNAM-3 at different wavelengths by the inhibition induced by OptoNAM-3 at 365 nm and in the dark since these two conditions represent respectively the minimum and maximum of the inhibition. We followed this equation: 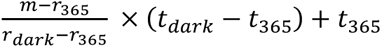 where *m* ∈ *[r*_365_*, r_dark_]* is the measurement (inhibition) to be scaled, *r*_365_ and *r_dark_* correspond respectively to the minimum (365 nm state inhibition) and maximum (dark state inhibition) of the measurement and *t*_365_ and *t_dark_* represent the minimum and maximum of the range (OptoNAM-3 365 nm PSS and dark PSS mean inhibition). To reduce the decreasing effect of normalization onto SEM, the SEM represented in the figures were calculated with the data injected in the first part of the equation 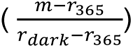.

To estimate the ratio of *trans*/*cis* isomers obtained upon illumination of bound OptoNAM-3 (Fig. 4E and J respectively), we calculated the residual currents obtained as described in the paragraph above and deduced their corresponding concentrations from the dose response curve equation of OptoNAM-3 in the dark (assuming dark state represents 100% *trans*). The obtained concentration was then divided by the initial *trans* concentration (2 µM) to obtain the percentage of *trans* isomer at each wavelength.

For the excitotoxicity test (Fig. S8), data were normalized by the mean of control (considered as 100% survival) and NMDA condition (considered as 0% survival) of the well plate. We followed this equation: 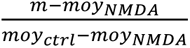 where *m* is the measurement (absorbance) to be normalized, *moy_NMDA_* and *moy_ctrl_* the mean absorbance of the NMDA and control condition for the well plate.

### Behavioral tests in Xenopus *laevis* tadpoles

Xenopus *laevis* embryos were obtained from the European Xenopus Resource Centre (EXRC, Portsmouth, UK) or TEFOR Paris Sackay (Saclay, France) and maintained in Modified Barth’s Saline (MBS) 0.1X (pH adjusted to 7.8 with NaOH), supplemented with 10 µg/mL of gentamicin, in 10 cm petri dishes with a maximum of 50 tadpoles per dish. The embryos were stored at 18°C until they reached stage 49 (12-14 days post-fertilization)(98).

For the experiments, (see Fig. S9A), stage 49 *X. laevis* tadpoles were placed in the center of a 12-well plate at a density of three animals per well in 0.1X MBS. The 12-well plate was covered with a 3D-printed dome equipped with a liquid guide (15 cm) connected to a multichannel LED delivery system (CoolLed pE-4000) at the top. The plate was positioned on optical cast plastic infrared filters, which allowed only wavelengths above 650 nm to pass, and red light (770 nm at 15% intensity) was continuously applied during recording to enable video recording in the dark. To habituate the tadpoles, the red light was gradually increased to 15% (0.03 mW) over a 15-minute period, and their baseline locomotion was recorded for 3 minutes. Subsequently, the tadpoles were incubated in the dark for 30 minutes in a 60 x 15 mm petri dish containing 5 mL of either the vehicle (0.1% DMSO in MBS 0.1X) or OptoNAM-3 solution (5 µM in MBS 0.1X) to facilitate drug penetration. Afterward, the tadpoles were transferred back to the 12-well plate, which contained either the vehicle solution or OptoNAM-3 (5 µM), and habituation to red light was performed again for 15 minutes. A three-minute video was then recorded to assess the locomotion in the dark after a 45-minute incubation and during cycles of one-minute illumination at 365 nm (or 460 nm) and 550 nm, alternating with three-minute rest periods (365 nm, 460 nm power ∼ 0.05 mW; 550 nm, power ∼ 0.36 mW). UV-green light cycles were consecutively repeated twice. Following the pharmacological treatments, the tadpoles were returned to water. The next day, the animals were observed for abnormalities.

To extract multi-animal tracking and pose estimation from the recorded tadpole videos, we employed the open-source deep learning toolbox Deeplabcut (73, 74). The toolbox is based on transfer learning and adapted from ImageNet-pretrained ResNets, specifically Resnet50, which is a model pretrained with over a million images. The original videos of the 12-well plates were cropped into individual wells to track the movement of three tadpoles per video. We selected 20 frames per video and labeled them with seven markers representing different body parts (right and left eye, stomach, and 4 points on the tail) to train the model for pose recognition using a multi-task convolutional neural network (CNN) that performs pose estimation by localizing key points in images. To enable the connection of key points to a given animal, additional deconvolution layers were added to the program (73). Model evaluation was conducted by calculating the pixel error between the predictions and manually labeled frames. Once the model achieved satisfactory training (network evaluation with a 1.3-pixel error for both training and testing), the video was analyzed, and individual tadpole traveled distances were calculated based on the x and y pixel coordinates of the labels provided by the data sheet obtained from video analysis. The stomach traveled distance yielded the best results; therefore, the tadpole traveled distance reported here represents the distance traveled by the "stomach" label. Since the identity of the 3 tadpoles could not be determined across the videos, the mean of the traveled distance per 3 tadpoles in one well was calculated in order to do a paired statistical test between light conditions. In the figures representing the *in vivo* experiment, one point therefore represents the mean locomotion of 3 tadpoles of the well. Tadpole locomotion was quantified by dividing the traveled distance by the recording duration and normalizing it twice: first by their baseline locomotion (see Fig S9 B, C) and secondly by the control (see Fig. 6).

### *In silico* docking and molecular dynamics

To model the protein/ligand interactions, we used the crystallographic structure 5EWJ (99). Missing residues were modeled by overlapping 5EWJ with the structure predicted by AlphaFold (100) for the same sequence. Residues 96 to 103 and 184 to 209 from chain A, and 42 to 65 from chain B were then extracted from the AlphaFold structure and added to the 5EWJ one to obtain a complete protein structure. Three-dimensional structures of the ligands Ifenprodil and OptoNAM-3 were obtained from their SMILES description with Gypsum-DL (101), and their protonation states were manually corrected. When four different starting conformations of Ifenprodil were docked in 5EWJ with Vina (102, 103), only one pose was found similar to the crystallographic conformation of ifenprodil within 5EWJ. However, when these conformations were docked with Gnina (104) (which is a fork of Vina that uses neural networks for the docking and the scoring), the four poses were found in a similar orientation as the experimental one. Thus, we have used Gnina as a docking engine. We add here that if the starting conformation of ifenprodil is the one extracted from 5EWJ (and not the ones coming from Gypsum-DL), the docked pose perfectly overlaps the crystallographic pose. The Ifenprodil pose with the best score (-11.2) is displayed in Figure S10A in green, together with the crystallographic pose that is displayed in orange. OptoNAM-3 was then docked: 10 different conformations were used, and the most relevant one (with the pyridinium of OptoNAM-3 overlapping with the phenol of ifenprodil, and the two terminal phenyls in the same pocket) is displayed in Figure S10B in purple (score of -9.5).

The geometry of OptoNAM-3 was then optimized at the M06-2X/6-31+G(d,p) level of geometry in a PCM of water with Gaussian09 A.02 (105). The RESP charges were then obtained at the HF/6-31G(d) level of geometry with Gaussian09 C.01 and acpype (106) and antechamber (107) were used to get the GAFF2 force field parameters for OptoNAM-3 (108). Finally, the force field was modified with an improved description of the azo moiety (109).

Molecular dynamics (MD) simulations were performed with Gromacs (104–106) with the Amber14SB force field to describe the protein atoms and ions (113) and the TIP3P description of water (114). After solvation in a rhombic odecahedron box with at least 8 Å between the solute atoms and the edges of the box, the system was neutralized with 18 sodium ions. The energy was then minimized with steepest descent to avoid steric clashes. The system was equilibrated during a 500 ps NPT simulation where we used a simulated annealing procedure to gradually heat the system from 100 K to 300 K in 400 ps; the system then evolved 100 ps at 300 K. During the equilibration, the velocity-rescale thermostat (115) and the Berendsen barostat (116) were used, with a time step of 1fs. Bonds containing an hydrogen were constrained with the LINCS algorithm (117, 118) with default parameters. Non-bonded interactions were described with PME for the electrostatics (119) with standard values for the force field (i.e. change between direct and reciprocal spaces at 8 Å) and default parameters, and cut-off of van der Waals interactions at 8 Å. We then performed the production simulation during 1 µs, where the parameters were similar to those in the equilibration step except for the barostat which was changed to Parrinello-Rahman (120) and for the time step which was increased to 2 fs.

### *In silico* quantum mechanics simulations

Quantum mechanics (QM) and quantum mechanics/molecular mechanics (QM/MM) calculations were performed using the ORCA 5.0.4 suite of programs (121). Geometry optimizations were carried out at the restricted Kohn-Sham Density Functional Theory (DFT) level without the use of symmetry, employing the PBE0 functional (122), the def2-TZVP basis set (123) with matching auxiliary basis sets (124) and the D3 correction (125, 126). The RIJCOSX approach applying the resolution of identity (RI) approximation to the Coulomb part and the chain of spheres (COS) seminumerical integration algorithm to the exchange term was used (127, 128). The convergence criteria for both the SCF was set to TIGHT and all the other parameters were chosen as default. For Time-Dependent DFT (TDDFT) calculations, the double-hybrid B2PLYP functional (129, 130) was chosen with at least 5 roots and without the TDA approximation. In more details, for calculations done in water the linear response conductor-like polarizable continuum model (LR-CPCM) perturbation of the density was included for both DFT and TDDFT calculations (131, 132). For calculations done in the protein the ORCA multiscale module was used. QM/MM calculations were performed using the additive scheme together with an electrostatic embedding. OptoNAM-3 consists of the QM level while the rest of the system is treated at the MM level using the AMBER14SB (108, 113, 133) force field as required by ORCA software. All the other parameters were chosen as default.

### Statistical analysis

Data are presented as mean ± standard deviation of the mean (SEM). All sample numbers (n) and statistical tests are specified in the figure legends. Statistical significances are indicated with *, ** and *** when p values are below 0.05, 0.01 and 0.001, respectively. n.s. indicates non-significant. Significance was defined as P < 0.05.

## Acknowledgments

We would like to thank Nicolas Delsuc (Chemistry department, ENS, Paris) for training and help on HPLC, Melissa David for help with molecular biology, Teddy Grand for help with electrophysiology, Julie Lefrançois and Cécile Cardoso for neuronal cultures and help with HEK cell culture, as well as Francine Acher for helpful discussions. This work was supported by PSL-QLife PhD fellowship (Q-life ANR-17-CONV-0005 PhD fellowship to C.G.), Labex Memolife (postdoc fellowship to C.G.), the European Commission (Marie-Sklodowska-Curie fellowship H2020-MSCA-IF-2015 Grant #701467 to L.M.) and the European Research Council (ERC Advanced Grant #693021 to P.P.).

## References

1. P. Paoletti, C. Bellone, Q. Zhou, NMDA receptor subunit diversity: impact on receptor properties, synaptic plasticity and disease. Nat Rev Neurosci 14, 383– 400 (2013).

2. K. B. Hansen, et al., Structure, Function, and Pharmacology of Glutamate Receptor Ion Channels. Pharmacol Rev 73, 1469–1658 (2021).

3. C. Geoffroy, P. Paoletti, L. Mony, Positive allosteric modulation of NMDA receptors: mechanisms, physiological impact and therapeutic potential. J Physiol (2021) 10.1113/JP280875.

4. J. W. Olney, Brain lesions, obesity, and other disturbances in mice treated with monosodium glutamate. Science 164, 719–721 (1969).

5. L. Mony, J. N. Kew, M. J. Gunthorpe, P. Paoletti, Allosteric modulators of NR2B-containing NMDA receptors: molecular mechanisms and therapeutic potential. British Journal of Pharmacology 157, 1301–1317 (2009).

6. C. G. Parsons, A. Stöffler, W. Danysz, Memantine: a NMDA receptor antagonist that improves memory by restoration of homeostasis in the glutamatergic system - too little activation is bad, too much is even worse. Neuropharmacology 53, 699–723 (2007).

7. J. Wang, F. Wang, D. Mai, S. Qu, Molecular Mechanisms of Glutamate Toxicity in Parkinson’s Disease. Front Neurosci 14, 585584 (2020).

8. G. E. Hardingham, H. Bading, Synaptic versus extrasynaptic NMDA receptor signalling: implications for neurodegenerative disorders. Nat Rev Neurosci 11, 682–696 (2010).

9. K. Gogas, Glutamate-based therapeutic approaches: NR2B receptor antagonists. Current Opinion in Pharmacology 6, 68–74 (2006).

10. D. Kreutzwiser, Q. A. Tawfic, Expanding Role of NMDA Receptor Antagonists in the Management of Pain. CNS Drugs 33, 347–374 (2019).

11. M. P. Parsons, L. A. Raymond, Extrasynaptic NMDA receptor involvement in central nervous system disorders. Neuron 82, 279–293 (2014).

12. S. Zhu, P. Paoletti, Allosteric modulators of NMDA receptors: multiple sites and mechanisms. Curr Opin Pharmacol 20, 14–23 (2015).

13. J. A. McCauley, NR2B subtype-selective NMDA receptor antagonists: 2001 – 2004. Expert Opinion on Therapeutic Patents 15, 389–407 (2005).

14. H. Ahmed, A. Haider, S. M. Ametamey, N-Methyl-D-Aspartate (NMDA) receptor modulators: a patent review (2015-present). Expert Opin Ther Pat 30, 743–767 (2020).

15. W. Liu, et al., A comprehensive description of GluN2B-selective N-methyl-D-aspartate (NMDA) receptor antagonists. European Journal of Medicinal Chemistry 200, 112447 (2020).

16. B. Gotti, et al., Ifenprodil and SL 82.0715 as cerebral anti-ischemic agents. I. Evidence for efficacy in models of focal cerebral ischemia. J Pharmacol Exp Ther 247, 1211–1221 (1988).

17. C. Carter, et al., Ifenprodil and SL 82.0715 as cerebral anti-ischemic agents. II. Evidence for N-methyl-D-aspartate receptor antagonist properties. J Pharmacol Exp Ther 247, 1222–1232 (1988).

18. K. Williams, Ifenprodil discriminates subtypes of the N-methyl-D-aspartate receptor: selectivity and mechanisms at recombinant heteromeric receptors. Mol Pharmacol 44, 851–859 (1993).

19. Y. Liu, et al., NMDA Receptor Subunits Have Differential Roles in Mediating Excitotoxic Neuronal Death Both In Vitro and In Vivo. J. Neurosci. 27, 2846– 2857 (2007).

20. J. von Engelhardt, et al., Excitotoxicity in vitro by NR2A- and NR2B-containing NMDA receptors. Neuropharmacology 53, 10–17 (2007).

21. H. Yuan, et al., Context-dependent GluN2B-selective inhibitors of NMDA receptor function are neuroprotective with minimal side effects. Neuron 85, 1305–1318 (2015).

22. C. Ikonomidou, L. Turski, Why did NMDA receptor antagonists fail clinical trials for stroke and traumatic brain injury? The Lancet Neurology 1, 383–386 (2002).

23. E. Karakas, N. Simorowski, H. Furukawa, Subunit arrangement and phenylethanolamine binding in GluN1/GluN2B NMDA receptors. Nature 475, 249–253 (2011).

24. D. Stroebel, et al., A Novel Binding Mode Reveals Two Distinct Classes of NMDA Receptor GluN2B-selective Antagonists. Mol Pharmacol 89, 541–551 (2016).

25. K. B. Hansen, et al., Structure, function, and allosteric modulation of NMDA receptors. Journal of General Physiology 150, 1081–1105 (2018).

26. N. Tajima, et al., Activation of NMDA receptors and the mechanism of inhibition by ifenprodil. Nature 534, 63–68 (2016).

27. L. Mony, S. Zhu, S. Carvalho, P. Paoletti, Molecular basis of positive allosteric modulation of GluN2B NMDA receptors by polyamines. EMBO J 30, 3134–3146 (2011).

28. J.-B. Esmenjaud, et al., An inter-dimer allosteric switch controls NMDA receptor activity. EMBO J 38 (2019).

29. M. Tian, et al., GluN2A and GluN2B NMDA receptors use distinct allosteric routes. Nat Commun 12, 4709 (2021).

30. M. Gielen, B. S. Retchless, L. Mony, J. W. Johnson, P. Paoletti, Mechanism of differential control of NMDA receptor activity by NR2 subunits. NATURE 459, 703–U107 (2009).

31. G. Fischer, et al., Ro 25-6981, a highly potent and selective blocker of N-methyl-D-aspartate receptors containing the NR2B subunit. Characterization in vitro. J Pharmacol Exp Ther 283, 1285–1292 (1997).

32. F. Menniti, et al., CP-101,606, a potent neuroprotectant selective for forebrain neurons. Eur J Pharmacol 331, 117–126 (1997).

33. B. L. Chenard, et al., (1S,2S)-1-(4-hydroxyphenyl)-2-(4-hydroxy-4-phenylpiperidino)-1-propanol: a potent new neuroprotectant which blocks N-methyl-D-aspartate responses. J Med Chem 38, 3138–3145 (1995).

34. F. Perin-Dureau, J. Rachline, J. Neyton, P. Paoletti, Mapping the Binding Site of the Neuroprotectant Ifenprodil on NMDA Receptors. J. Neurosci. 22, 5955–5965 (2002).

35. P. Legendre, G. L. Westbrook, Ifenprodil blocks N-methyl-D-aspartate receptors by a two-component mechanism. Mol Pharmacol 40, 289–298 (1991).

36. J. N. Kew, J. A. Kemp, An allosteric interaction between the NMDA receptor polyamine and ifenprodil sites in rat cultured cortical neurones. J Physiol 512 **(Pt** **1**), 17–28 (1998).

37. J. N. Kew, G. Trube, J. A. Kemp, State-dependent NMDA receptor antagonism by Ro 8-4304, a novel NR2B selective, non-competitive, voltage-independent antagonist. Br J Pharmacol 123, 463–472 (1998).

38. C. Bellone, R. A. Nicoll, Rapid bidirectional switching of synaptic NMDA receptors. Neuron 55, 779–785 (2007).

39. T. Papouin, et al., Synaptic and extrasynaptic NMDA receptors are gated by different endogenous coagonists. Cell 150, 633–646 (2012).

40. P. Paoletti, G. C. R. Ellis-Davies, A. Mourot, Optical control of neuronal ion channels and receptors. Nat Rev Neurosci 20, 514–532 (2019).

41. R. H. Kramer, A. Mourot, H. Adesnik, Optogenetic pharmacology for control of native neuronal signaling proteins. Nat Neurosci 16, 816–823 (2013).

42. K. Hüll, J. Morstein, D. Trauner, In Vivo Photopharmacology. Chem. Rev. 118, 10710–10747 (2018).

43. E. Merino, M. Ribagorda, Control over molecular motion using the *cis* – *trans* photoisomerization of the azo group. Beilstein J. Org. Chem. 8, 1071–1090 (2012).

44. E. R. Thapaliya, et al., Photochemical Control of Drug Efficacy: A Comparison of Uncaging and Photoswitching Ifenprodil on NMDA Receptors. ChemPhotoChem 5, 445–454 (2021).

45. L. Mony, N. Triballeau, P. Paoletti, F. C. Acher, H.-O. Bertrand, Identification of a novel NR2B-selective NMDA receptor antagonist using a virtual screening approach. Bioorg Med Chem Lett 20, 5552–5558 (2010).

46. J. Broichhagen, J. A. Frank, D. Trauner, A Roadmap to Success in Photopharmacology. Acc. Chem. Res. 48, 1947–1960 (2015).

47. J. Morstein, M. Awale, J.-L. Reymond, D. Trauner, Mapping the Azolog Space Enables the Optical Control of New Biological Targets. ACS Cent Sci 5, 607– 618 (2019).

48. W. A. Velema, W. Szymanski, B. L. Feringa, Photopharmacology: Beyond Proof of Principle. J. Am. Chem. Soc. 136, 2178–2191 (2014).

49. X. Gómez-Santacana, et al., Illuminating Phenylazopyridines To Photoswitch Metabotropic Glutamate Receptors: From the Flask to the Animals. ACS Cent. Sci. 3, 81–91 (2017).

50. S. Pittolo, et al., An allosteric modulator to control endogenous G protein-coupled receptors with light. Nat Chem Biol 10, 813–815 (2014).

51. C. Zussy, et al., Dynamic modulation of inflammatory pain-related affective and sensory symptoms by optical control of amygdala metabotropic glutamate receptor 4. Mol Psychiatry 23, 509–520 (2018).

52. A. Landra-Willm, et al., A photoswitchable inhibitor of TREK channels controls pain in wild-type intact freely moving animals. Nat Commun 14, 1160 (2023).

53. N. J. Liverton, et al., Identification and characterization of 4-methylbenzyl 4-[(pyrimidin-2-ylamino)methyl]piperidine-1-carboxylate, an orally bioavailable, brain penetrant NR2B selective N-methyl-D-aspartate receptor antagonist. J Med Chem 50, 807–819 (2007).

54. B. Büttelmann, et al., 4-(3,4-Dihydro-1H-isoquinolin-2yl)-pyridines and 4-(3,4-Dihydro-1H-isoquinolin-2-yl)-quinolines as potent NR1/2B subtype selective NMDA receptor antagonists. Bioorganic & Medicinal Chemistry Letters 13, 1759–1762 (2003).

55. J. A. McCauley, et al., NR2B-Selective *N* -Methyl-D-aspartate Antagonists: Synthesis and Evaluation of 5-Substituted Benzimidazoles. J. Med. Chem. 47, 2089–2096 (2004).

56. 56. A. Alanine, B. Buettelmann, M.-P. H. Neidhart, E. Pinard, R. Wyler, Pyridine derivatives as nmda-receptor subtype blockers (2003) (April 19, 2022).

57. D. Stroebel, S. Carvalho, T. Grand, S. Zhu, P. Paoletti, Controlling NMDA Receptor Subunit Composition Using Ectopic Retention Signals. J. Neurosci. 34, 16630–16636 (2014).

58. K. B. Hansen, K. K. Ogden, H. Yuan, S. F. Traynelis, Distinct functional and pharmacological properties of Triheteromeric GluN1/GluN2A/GluN2B NMDA receptors. Neuron 81, 1084–1096 (2014).

59. J. Rachline, The Micromolar Zinc-Binding Domain on the NMDA Receptor Subunit NR2B. Journal of Neuroscience 25, 308–317 (2005).

60. L. Mony, et al., Structural basis of NR2B-selective antagonist recognition by N-methyl-D-aspartate receptors. Mol Pharmacol 75, 60–74 (2009).

61. J. A. Kemp, T. Tasker, Methods for treating disorders using nmda nr2b-subtype selective antagonist (2010) (March 7, 2022).

62. S. McKay, et al., The Developmental Shift of NMDA Receptor Composition Proceeds Independently of GluN2 Subunit-Specific GluN2 C-Terminal Sequences. Cell Reports 25, 841–851.e4 (2018).

63. K. Williams, S. L. Russell, Y. M. Shen, P. B. Molinoff, Developmental switch in the expression of NMDA receptors occurs in vivo and in vitro. Neuron 10, 267– 278 (1993).

64. M. Homocianu, Optical properties of solute molecules: Environmental effects, challenges, and their practical implications. Microchemical Journal 161, 105797 (2021).

65. A. B. Grommet, L. M. Lee, R. Klajn, Molecular Photoswitching in Confined Spaces. Acc. Chem. Res. 53, 2600–2610 (2020).

66. J. von Engelhardt, et al., Excitotoxicity in vitro by NR2A- and NR2B-containing NMDA receptors. Neuropharmacology 53, 10–17 (2007).

67. K. Sillar, wen-chang li, “Neural control of swimming in hatchling Xenopus frog tadpoles” in (2020), pp. 153–174.

68. W.-C. Li, A. Roberts, S. R. Soffe, Specific Brainstem Neurons Switch Each Other into Pacemaker Mode to Drive Movement by Activating NMDA Receptors. J. Neurosci. 30, 16609–16620 (2010).

69. J. P. Issberner, K. T. Sillar, The contribution of the NMDA receptor glycine site to rhythm generation during fictive swimming in Xenopus laevis tadpoles. Eur J Neurosci 26, 2556–2564 (2007).

70. R. Ewald, H. Cline, Cloning and phylogenetic analysis of NMDA receptor subunits NR1, NR2A and NR2B in Xenopus laevis tadpoles. Frontiers in Molecular Neuroscience 2 (2009).

71. C. Schmidt, M. Hollmann, Apparent Homomeric NR1 Currents Observed in Xenopus Oocytes are Caused by an Endogenous NR2 Subunit. Journal of Molecular Biology 376, 658–670 (2008).

72. H.-Y. Zhang, L. Picton, W.-C. Li, K. T. Sillar, Mechanisms underlying the activity-dependent regulation of locomotor network performance by the Na+ pump. Sci Rep 5, 16188 (2015).

73. J. Lauer, et al., “Multi-animal pose estimation and tracking with DeepLabCut” (Animal Behavior and Cognition, 2021) 10.1101/2021.04.30.442096 (January 23, 2022).

74. A. Mathis, et al., DeepLabCut: markerless pose estimation of user-defined body parts with deep learning. Nat Neurosci 21, 1281–1289 (2018).

75. S. P. Currie, G. H. Doherty, K. T. Sillar, Deep-brain photoreception links luminance detection to motor output in Xenopus frog tadpoles. Proc Natl Acad Sci U S A 113, 6053–6058 (2016).

76. F. Perin-Dureau, J. Rachline, J. Neyton, P. Paoletti, Mapping the binding site of the neuroprotectant ifenprodil on NMDA receptors. J Neurosci 22, 5955–5965 (2002).

77. M. Volgraf, et al., Allosteric control of an ionotropic glutamate receptor with an optical switch. Nat Chem Biol 2, 47–52 (2006).

78. D. B. Konrad, et al., Computational Design and Synthesis of a Deeply Red-Shifted and Bistable Azobenzene. J. Am. Chem. Soc. 142, 6538–6547 (2020).

79. C. L. Forber, E. C. Kelusky, N. J. Bunce, M. C. Zerner, Electronic spectra of cis- and trans-azobenzenes: consequences of ortho substitution. J. Am. Chem. Soc. 107, 5884–5890 (1985).

80. A. A. Beharry, O. Sadovski, G. A. Woolley, Azobenzene Photoswitching without Ultraviolet Light. J. Am. Chem. Soc. 133, 19684–19687 (2011).

81. D. Bléger, J. Schwarz, A. M. Brouwer, S. Hecht, o-Fluoroazobenzenes as Readily Synthesized Photoswitches Offering Nearly Quantitative Two-Way Isomerization with Visible Light. J. Am. Chem. Soc. 134, 20597–20600 (2012).

82. D. M. Barber, et al., Optical control of AMPA receptors using a photoswitchable quinoxaline-2,3-dione antagonist. Chem. Sci. 8, 611–615 (2016).

83. M. Ricart-Ortega, J. Font, A. Llebaria, GPCR photopharmacology. Molecular and Cellular Endocrinology 488, 36–51 (2019).

84. J. A. R. Dalton, et al., Shining Light on an mGlu5 Photoswitchable NAM: A Theoretical Perspective. Curr Neuropharmacol 14, 441–454 (2016).

85. L. Laprell, et al., Optical control of NMDA receptors with a diffusible photoswitch. Nat Commun 6, 8076 (2015).

86. M. Volgraf, et al., Reversibly Caged Glutamate: A Photochromic Agonist of Ionotropic Glutamate Receptors. J. Am. Chem. Soc. 129, 260–261 (2007).

87. F. W. W. Hartrampf, et al., Development of a photoswitchable antagonist of NMDA receptors. Tetrahedron 73, 4905–4912 (2017).

88. P. Stawski, M. Sumser, D. Trauner, A Photochromic Agonist of AMPA Receptors. Angewandte Chemie International Edition 51, 5748–5751 (2012).

89. M. Nikolaev, D. Tikhonov, Light-Sensitive Open Channel Block of Ionotropic Glutamate Receptors by Quaternary Ammonium Azobenzene Derivatives. Int J Mol Sci 24, 13773 (2023).

90. M. V. Nikolaev, D. M. Strashkov, M. N. Ryazantsev, D. B. Tikhonov, Development of a quaternary ammonium photoswitchable antagonist of NMDA receptors. Eur J Pharmacol 938, 175448 (2023).

91. P. J. Conn, A. Christopoulos, C. W. Lindsley, Allosteric modulators of GPCRs: a novel approach for the treatment of CNS disorders. Nat Rev Drug Discov 8, 41– 54 (2009).

92. J.-P. Changeux, A. Christopoulos, Allosteric Modulation as a Unifying Mechanism for Receptor Function and Regulation. Cell 166, 1084–1102 (2016).

93. C. J. Hatton, P. Paoletti, Modulation of triheteromeric NMDA receptors by N-terminal domain ligands. Neuron 46, 261–74 (2005).

94. P. Paoletti, J. Neyton, P. Ascher, Glycine-independent and subunit-specific potentiation of NMDA responses by extracellular Mg2+. Neuron 15, 1109–1120 (1995).

95. P. Paoletti, P. Ascher, J. Neyton, High-affinity zinc inhibition of NMDA NR1-NR2A receptors. J Neurosci 17, 5711–5725 (1997).

96. M. L. Seibenhener, M. W. Wooten, Isolation and Culture of Hippocampal Neurons from Prenatal Mice. J Vis Exp, 3634 (2012).

97. T. Mosmann, Rapid colorimetric assay for cellular growth and survival: Application to proliferation and cytotoxicity assays. Journal of Immunological Methods 65, 55–63 (1983).

98. H. L. Sive, R. M. Grainger, R. M. Harland, Housing and Feeding of Xenopus laevis. CSH Protoc 2007, pdb.top8 (2007).

99. D. Stroebel, et al., A Novel Binding Mode Reveals Two Distinct Classes of NMDA Receptor GluN2B-selective Antagonists. Mol Pharmacol 89, 541–551 (2016).

100. J. Jumper, et al., Highly accurate protein structure prediction with AlphaFold. Nature 596, 583–589 (2021).

101. P. J. Ropp, et al., Gypsum-DL: an open-source program for preparing small-molecule libraries for structure-based virtual screening. Journal of Cheminformatics 11, 34 (2019).

102. O. Trott, A. J. Olson, AutoDock Vina: improving the speed and accuracy of docking with a new scoring function, efficient optimization, and multithreading. J Comput Chem 31, 455–461 (2010).

103. J. Eberhardt, D. Santos-Martins, A. F. Tillack, S. Forli, AutoDock Vina 1.2.0: New Docking Methods, Expanded Force Field, and Python Bindings. J Chem Inf Model 61, 3891–3898 (2021).

104. A. T. McNutt, et al., GNINA 1.0: molecular docking with deep learning. Journal of Cheminformatics 13, 43 (2021).

105. M. Frisch, et al., Gaussian 09 (Revision A02). Gaussian Inc. Wallingford CT (2009).

106. A. W. Sousa da Silva, W. F. Vranken, ACPYPE - AnteChamber PYthon Parser interfacE. BMC Research Notes 5, 367 (2012).

107. J. Wang, W. Wang, P. A. Kollman, D. A. Case, Automatic atom type and bond type perception in molecular mechanical calculations. J Mol Graph Model 25, 247–260 (2006).

108. J. Wang, R. M. Wolf, J. W. Caldwell, P. A. Kollman, D. A. Case, Development and testing of a general amber force field. Journal of Computational Chemistry 25, 1157–1174 (2004).

109. P. Duchstein, C. Neiss, A. Görling, D. Zahn, Molecular mechanics modeling of azobenzene-based photoswitches. J Mol Model 18, 2479–2482 (2012).

110. B. Hess, C. Kutzner, D. van der Spoel, E. Lindahl, GROMACS 4: Algorithms for Highly Efficient, Load-Balanced, and Scalable Molecular Simulation. J. Chem. Theory Comput. 4, 435–447 (2008).

111. S. Pronk, et al., GROMACS 4.5: a high-throughput and highly parallel open source molecular simulation toolkit. Bioinformatics 29, 845–854 (2013).

112. M. J. Abraham, et al., GROMACS: High performance molecular simulations through multi-level parallelism from laptops to supercomputers. SoftwareX 1–2, 19–25 (2015).

113. J. A. Maier, et al., ff14SB: Improving the Accuracy of Protein Side Chain and Backbone Parameters from ff99SB. J Chem Theory Comput 11, 3696–3713 (2015).

114. W. L. Jorgensen, Quantum and statistical mechanical studies of liquids. 10. Transferable intermolecular potential functions for water, alcohols, and ethers. Application to liquid water. ACS Publications (1981) 10.1021/ja00392a016 (September 27, 2023).

115. G. Bussi, T. Zykova-Timan, M. Parrinello, Isothermal-isobaric molecular dynamics using stochastic velocity rescaling. The Journal of Chemical Physics 130, 074101 (2009).

116. H. J. C. Berendsen, J. P. M. Postma, W. F. Van Gunsteren, A. DiNola, J. R. Haak, Molecular dynamics with coupling to an external bath. The Journal of Chemical Physics 81, 3684–3690 (1984).

117. B. Hess, P-LINCS: A Parallel Linear Constraint Solver for Molecular Simulation. J. Chem. Theory Comput. 4, 116–122 (2008).

118. B. Hess, H. Bekker, H. J. C. Berendsen, J. G. E. M. Fraaije, LINCS: A linear constraint solver for molecular simulations. Journal of Computational Chemistry 18, 1463–1472 (1997).

119. T. Darden, D. York, L. Pedersen, Particle mesh Ewald: An Nṡlog(N) method for Ewald sums in large systems. Journal of Chemical Physics 98, 10089–10092 (1993).

120. M. Parrinello, A. Rahman, Polymorphic transitions in single crystals: A new molecular dynamics method. Journal of Applied Physics 52, 7182–7190 (1981).

121. F. Neese, Software update: The ORCA program system—Version 5.0. WIREs Computational Molecular Science 12, e1606 (2022).

122. C. Adamo, V. Barone, Toward reliable density functional methods without adjustable parameters: The PBE0 model. The Journal of Chemical Physics 110, 6158–6170 (1999).

123. F. Weigend, R. Ahlrichs, Balanced basis sets of split valence, triple zeta valence and quadruple zeta valence quality for H to Rn: Design and assessment of accuracy. Phys. Chem. Chem. Phys. 7, 3297–3305 (2005).

124. F. Weigend, Accurate Coulomb-fitting basis sets for H to Rn. Phys. Chem. Chem. Phys. 8, 1057–1065 (2006).

125. S. Grimme, J. Antony, S. Ehrlich, H. Krieg, A consistent and accurate ab initio parametrization of density functional dispersion correction (DFT-D) for the 94 elements H-Pu. J Chem Phys 132, 154104 (2010).

126. A. D. Becke, E. R. Johnson, A density-functional model of the dispersion interaction. J Chem Phys 123, 154101 (2005).

127. B. Helmich-Paris, B. de Souza, F. Neese, R. Izsák, An improved chain of spheres for exchange algorithm. The Journal of Chemical Physics 155, 104109 (2021).

128. R. Izsák, F. Neese, An overlap fitted chain of spheres exchange method. The Journal of Chemical Physics 135, 144105 (2011).

129. S. Grimme, Semiempirical hybrid density functional with perturbative second-order correlation. J Chem Phys 124, 034108 (2006).

130. S. Grimme, F. Neese, Double-hybrid density functional theory for excited electronic states of molecules. The Journal of Chemical Physics 127, 154116 (2007).

131. V. Barone, M. Cossi, Quantum Calculation of Molecular Energies and Energy Gradients in Solution by a Conductor Solvent Model. J. Phys. Chem. A 102, 1995–2001 (1998).

132. M. Garcia-Ratés, F. Neese, Effect of the Solute Cavity on the Solvation Energy and its Derivatives within the Framework of the Gaussian Charge Scheme. J Comput Chem 41, 922–939 (2020).

133. D.A. Case, H.M. Aktulga, K. Belfon, I.Y. Ben-Shalom, J.T. Berryman, S.R. Brozell, D.S. Cerutti, T.E. Cheatham, III, G.A. Cisneros, V.W.D. Cruzeiro, T.A. Darden, N. Forouzesh, G. Giambaşu, T. Giese, M.K. Gilson, H. Gohlke, A.W. Goetz, J. Harris, S. Izadi, S.A. Izmailov, K. Kasavajhala, M.C. Kaymak, E. King, A. Kovalenko, T. Kurtzman, T.S. Lee, P. Li, C. Lin, J. Liu, T. Luchko, R. Luo, M. Machado, V. Man, M. Manathunga, K.M. Merz, Y. Miao, O. Mikhailovskii, G. Monard, H. Nguyen, K.A. O’Hearn, A. Onufriev, F. Pan, S. Pantano, R. Qi, A. Rahnamoun, D.R. Roe, A. Roitberg, C. Sagui, S. Schott-Verdugo, A. Shajan, J. Shen, C.L. Simmerling, N.R. Skrynnikov, J. Smith, J. Swails, R.C. Walker, J. Wang, J. Wang, H. Wei, X. Wu, Y. Wu, Y. Xiong, Y. Xue, D.M. York, S. Zhao, Q. Zhu, and P.A. Kollman, Amber 2023 (2023).

