## Supplementary information for "Reversible inhibition of GluN2B-containing NMDA receptors with an *in situ* red-shifted, photoswitchable antagonist"

**This PDF file includes:**

Supporting text
Figures S1 to S10
Tables S1 to S4
SI References

### Supporting Text

#### Photochemical and biological characterizations of OptoNAM-1, -2 and -4.

To characterize the photochemical properties of OptoNAM-1, -2 and -4, we acquired UV-visible spectra of these compounds in the dark (*trans* state) and after illumination with various wavelengths. The absorption spectra of OptoNAM-1, -2 and 4 in the dark were characteristic of azobenzenes in their *trans* configuration (black curves in Fig. S1B, F and J) (1). Application of UV light at a wavelength close to the main absorption peak of the *trans* form (365 nm) gave a completely different spectrum (violet curves in Fig. S1B, F and J), characteristic of azobenzenes in their *cis* configuration (1).

To determine the appropriate wavelengths to convert the *cis* OptoNAMs back to *trans*, we illuminated the 365 nm PSS OptoNAM-1, -2 and -4 solutions with wavelengths of 440 and 490 nm. 440 nm irradiation provided the strongest *cis*-to-*trans* conversion for OptoNAM-1 and -4 (OptoNAM-1, ~75% *trans* measured at $\lambda_{trans}$ = 320 nm; OptoNAM-4, 100% *trans* measured at $\lambda_{trans}$ = 334 nm), while 490 nm was most efficient to convert cis-OptoNAM-2 back to *trans* (~52% measured at $\lambda_{trans}$ = 371 nm) (Fig. S1A, B, E, F, I, J). The cis isomer of all compounds displayed strong photostability since the absorption spectrum of the 365 nm PSS kept in the dark did not change over 24h (Fig. S1C, G, K).

The activities of the dark and 365 nm PSS of OptoNAM-1, -2 and -4 on GluN1/GluN2B NMDARs were assessed by electrophysiology on Xenopus oocytes as described in the main text for OptoNAM-3. OptoNAM-1 and -2 behaved as GluN1/GluN2B NAMs with better apparent affinity in the dark than in the UV condition. However, OptoNAM-1 and 2 in *trans* configuration displayed a drastic loss of potency compared to their parent compounds (2–4), with a >1000-fold shift in potency (Fig. S1D, H and Table S1). We showed that replacement of the amino-methyl bond of the parent compounds by an azo bond to obtain OptoNAM-1 and -2 induced a loss of protonation of the aminopyridium moiety at physiological pH, which likely disrupts binding of the compounds in the ifenprodil binding site (Fig. S2A, B, D). OptoNAM-4, on the other hand, did not display any photodependence of activity (Fig. S1L and Table S1).

### Supplementary Figures

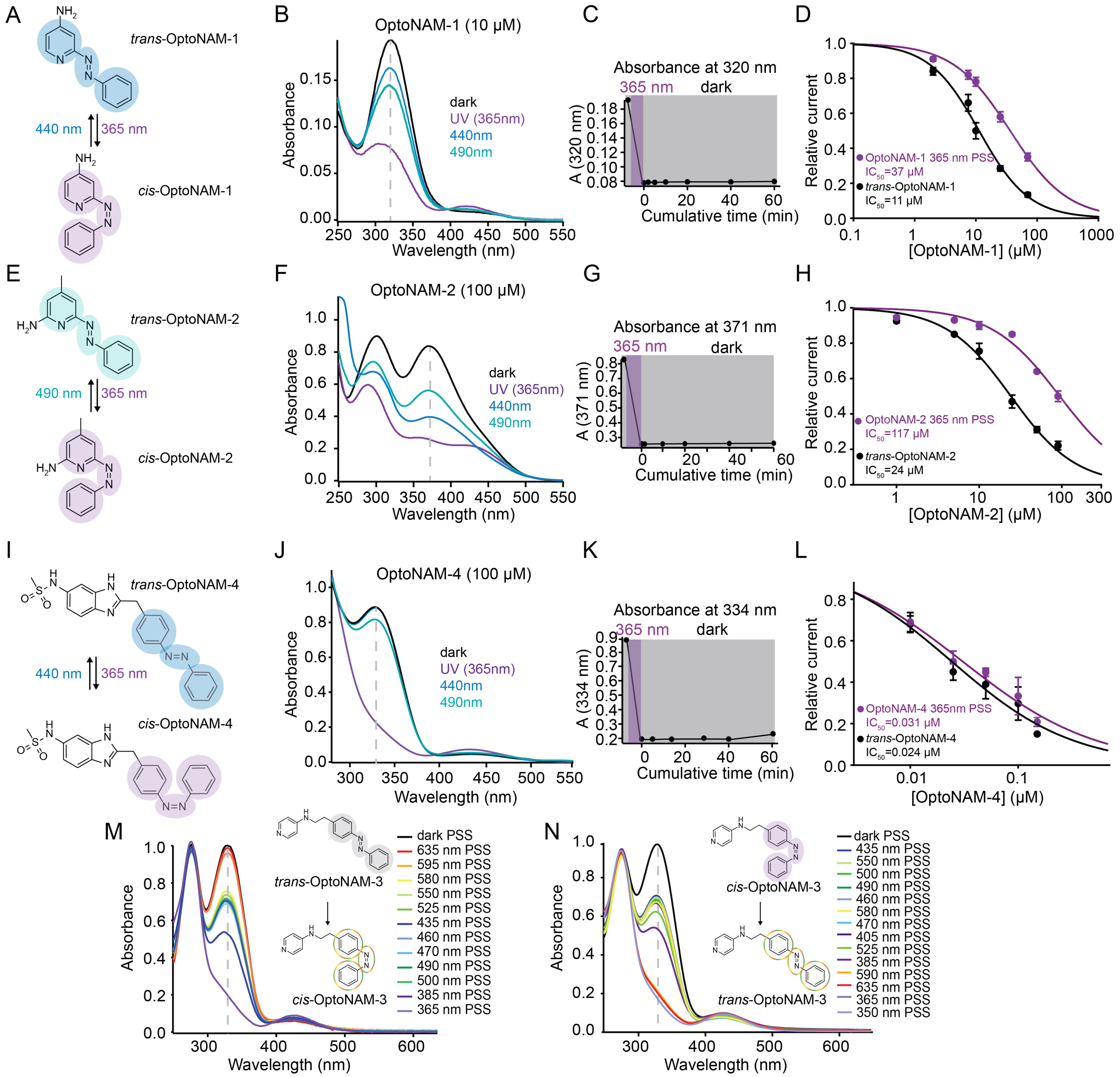

**Figure S1. Photochemical properties of OptoNAM-1 to -4 and their photodependent activity at GluN1/GluN2B receptors**

**(A-D) OptoNAM-1. (A)** In solution, OptoNAM-1 can be switched from *trans* to *cis* configuration by UV illumination (365 nm) and back to *trans* by 440 nm light. **(B)** UV-visible absorption spectra of OptoNAM-1 in the dark (black curve), after 365 nm illumination (violet curve) and subsequent illumination of the 365 nm PSS by 440 or 490 nm light. Dashed line represents the wavelength of peak absorption of *trans*-OptoNAM-1 (320 nm). **(C)** OptoNAM-1 365 nm PSS (mostly *cis*) is highly photostable in the dark, no change of the absorption spectra was observed up to 60 min after 365 nm illumination. **(D)** Dose-response curves of OptoNAM-1 activity on GluN1/GluN2B receptors in the dark (black curve, $IC_{50}$ = 11 ± 1 μM, n= 4-6) or pre-illuminated with 365 nm (violet curve, $IC_{50}$ = 37 ± 2 μM, n= 5-17).

**(E-H) OptoNAM-2. (E)** In solution, OptoNAM-2 can be switched from *trans* to *cis* by UV illumination (365 nm) and back to *trans* by 490 nm light. **(F)** UV-visible absorption spectra of OptoNAM-2 in the dark (black curve), after 365 nm illumination (violet curve) and subsequent illumination of the 365 nm PSS by 440 or 490 nm light. Dashed line represents the wavelength of peak absorption of *trans*-OptoNAM-2 (371 nm). **(G)** OptoNAM-2 365 nm PSS (mostly *cis*) is highly photostable in the dark, no change of the absorption spectra was observed up to 60 min after 365 nm illumination. **(H)** Dose-response curves of OptoNAM-2 activity on GluN1/GluN2B receptors in the dark (black curve, $IC_{50}$ = 24 ± 2 μM, n= 3-5) or pre-illuminated with 365 nm (violet curve, $IC_{50}$ = 117 ± 10 μM, n= 3-8).

**(I-L) OptoNAM-4. (I)** In solution, OptoNAM-4 can be switched from *trans* to *cis* by UV illumination (365 nm) and back to *trans* by 440 nm light. **(J)** UV-visible absorption spectra of OptoNAM-4 in the dark (black curve), after 365 nm illumination (violet curve) and subsequent illumination of the 365 nm PSS by 440 or 490 nm light. Note that 440 nm allows full return to the dark state. Dashed line represents the wavelength of peak absorption of *trans*-OptoNAM-2 (334 nm). **(K)** OptoNAM-4 365 nm PSS (mostly *cis*) is highly photostable in the dark, only little change of the absorption spectra was observed up to 60 min after 365 nm illumination. **(L)** Dose-response curves of OptoNAM-4 activity on GluN1/GluN2B receptors in the dark (black curve, $IC_{50}$ = 24 ± 26 nM, n = 3) or pre-illuminated with 365 nm (violet curve, $IC_{50}$ = 31 ± 29 nM, n = 3).

**(M-N) OptoNAM-3 *trans-cis* and *cis-trans* isomerization in solution. (M)** OptoNAM-3 UV-visible absorption spectra in the dark (black curve) and PSS obtained after illumination with wavelengths ranging from 350 to 635 nm of the dark PSS. These spectra were used to create panel D and E from Figure 4. **(N)** OptoNAM-3 UV-visible absorption spectra in the dark (black curve), after 365 nm illumination (violet curve), and PSS obtained after illumination with wavelengths ranging from 350 to 635 nm of the 365 nm PSS. These spectra were used to create panels I and J from Figure 4.

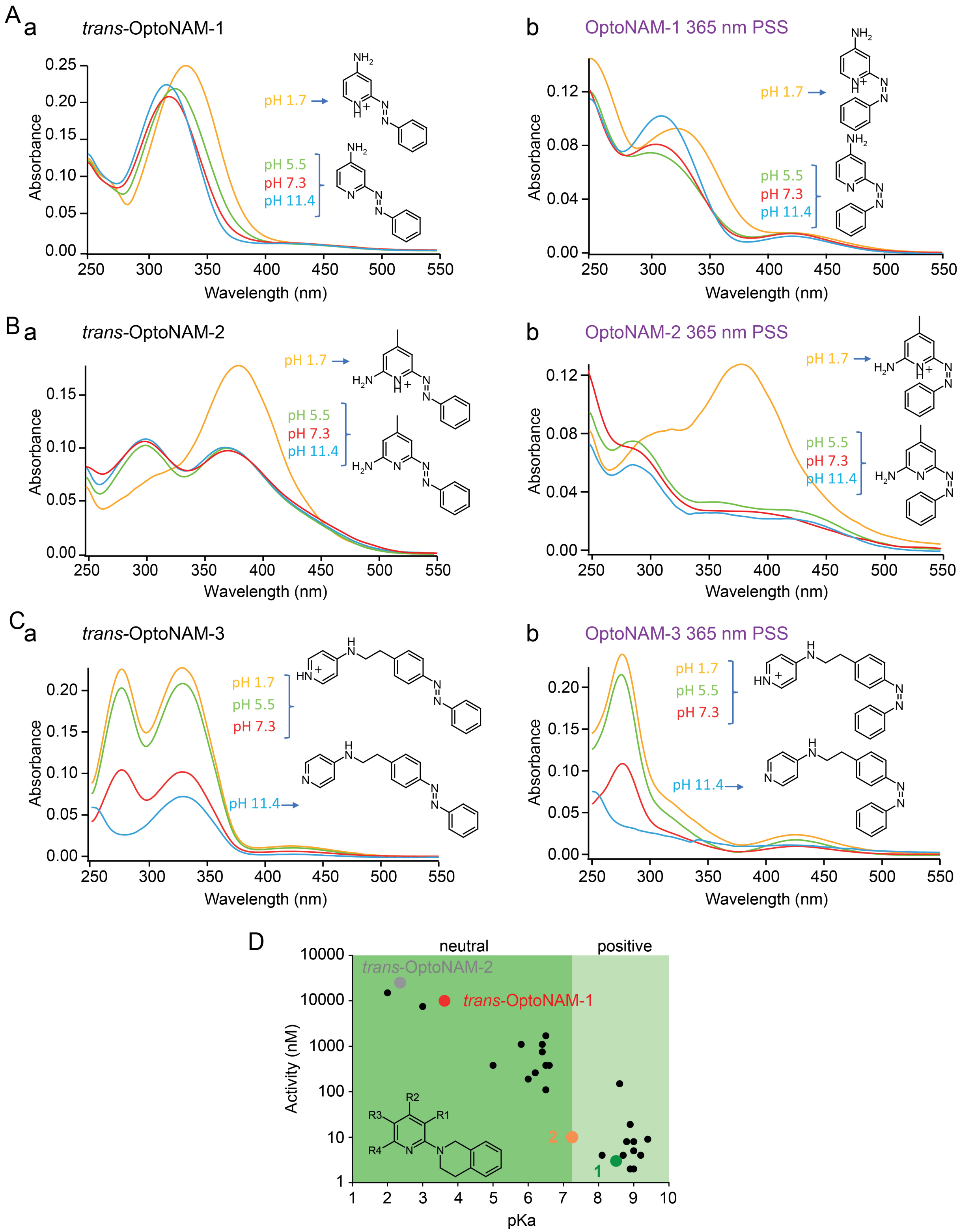
A a
trans-OptoNAM-1
Absorbance
Wavelength (nm)
pH 1.7
pH 5.5
pH 7.3
pH 11.4
b
OptoNAM-1 365 nm PSS
B a
trans-OptoNAM-2
b
OptoNAM-2 365 nm PSS
C a
trans-OptoNAM-3
b
OptoNAM-3 365 nm PSS
D
neutral
positive
trans-OptoNAM-2
trans-OptoNAM-1
Activity (nM)
pKa
R1
R2
R3
R4
1
2

**Figure S2. Decreased pKa of OptoNAM-1 and -2 compared to their parent compounds are likely responsible for their decreased activity. (A-C)** pKa estimation of the *trans* (a) and *cis* (365 nm PSS, b) isomers of OptoNAM-1 (A), -2 (B) and -3 (C) by UV-visible spectrum analysis. UV-visible absorption spectra were measured at different pH (1.7 in yellow, 5.5 in green, 7.3 in red and 11.4 in blue). Correspondence between spectra at different pH and OptoNAM protonated and unprotonated chemical structures is indicated. A leftward shift and a decrease of the absorbance peak corresponding to the aminopyridine carrying the charge was observed upon OptoNAM deprotonation. This effect is most obvious for OptoNAM-2 and characteristic of the deprotonated spectra of 2,6 diaminopyridine bases (5). Based on this analysis, at physiological pH, OptoNAM-1 and -2, either in *cis* or *trans*, are unprotonated, while OptoNAM-3 is protonated. **(D)** Relationship between the activity and the measured pKa of compounds from the same chemical series as parent compound **1** (values from ref. (6)) (black dots). Parent compound **1** is highlighted as a thick green dot. Parent compound **2** (orange), as well as *trans*-OptoNAM-1 (red) and -2 (grey) were added to the plot according to their published or measured activity (Table S1), and predicted pKa (pKa was predicted by Marvin, Chemaxon https://www.chemaxon.com). Note the tight correlation between pKa and activity, suggesting that the decreased pKa of OptoNAM-1 and -2 induced by azologization of the parent compounds, resulting in a loss of protonation at physiological pH, is responsible for the large decrease of activity of these compounds. “Neutral” (dark green) and “Positive” (light green) indicate the pKa ranges for which the compounds are neutral and positively-charged, respectively, at physiological pH (pH = 7.3).

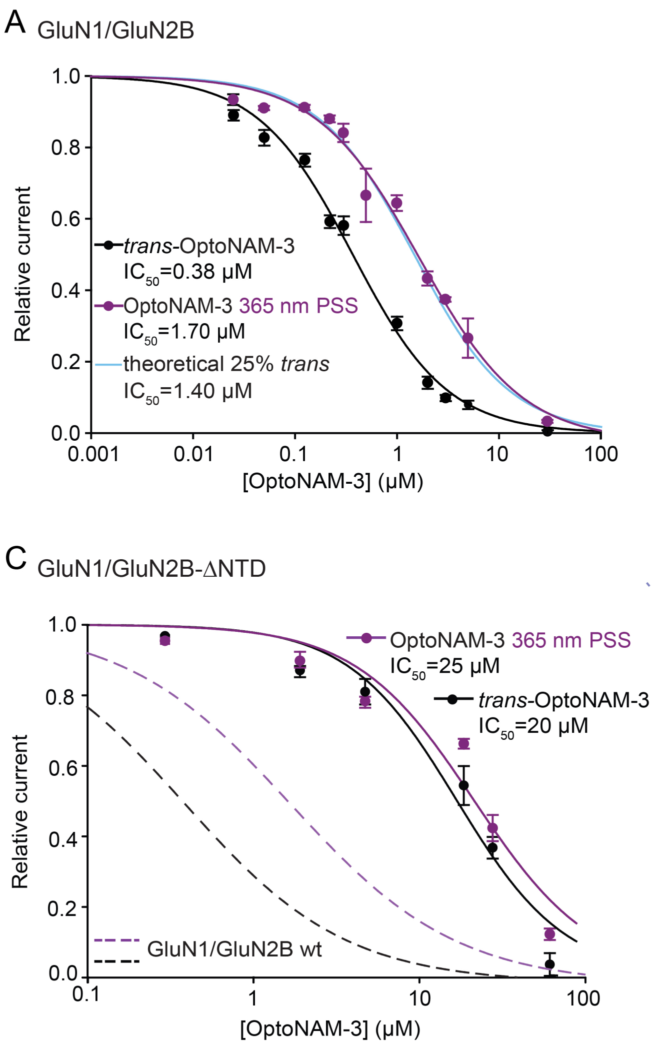
A GluN1/GluN2B
Relative current
[OptoNAM-3] (μM)
trans-OptoNAM-3
IC50=0.38 μM
OptoNAM-3 365 nm PSS
IC50=1.70 μM
theoretical 25% trans
IC50=1.40 μM
B GluN1/GluN2B
trans-OptoNAM-3
IC50=0.38 μM
OptoNAM-3 350 nm PSS
IC50=4.38 μM
theoretical 6% trans
IC50=6.15 μM
C GluN1/GluN2B-ΔNTD
OptoNAM-3 365 nm PSS
IC50=25 μM
trans-OptoNAM-3
IC50=20 μM
GluN1/GluN2B wt
D GluN1/GluN2B
+ ifenprodil
IC50=1.0 μM
- ifenprodil
IC50=0.38 μM

**Figure S3. Additional data relative to Figure 2.**

**(A) Inhibition by OptoNAM-3 365 nm PSS is exclusively mediated by the remaining *trans* isomer still present in solution.** Dose-response curves of OptoNAM-3 activity on GluN1/GluN2B in the dark (black curve, $IC_{50}$ = 0.38 ± 0.03 µM, n= 5-21), pre-illuminated by 365 nm (purple curve, $IC_{50}$ = 1.70 ± 0.20 µM, n= 5-17) and theoretical dose-response curve (blue) of a mixture of 25% *trans*- and 75% *cis*-OptoNAM-3 (corresponding to the 365 nm PSS determined by HPLC) assuming that only the *trans* isomer is active. The theoretical curve was calculated from the *trans* dose-response curve in the dark. The 25% *trans* theoretical dose-response curve is perfectly superimposable to the one of OptoNAM-3 365 nm PSS. (**B)** Dose-response curves of OptoNAM-3 activity on GluN1/GluN2B in the dark (black curve, $IC_{50}$ = 0.38 ± 0.03 µM, n= 5-21), pre-illuminated by 350 nm (lavender curve, $IC_{50}$ = 4.38 ± 0.59 µM n= 4-13) and theoretical dose-response curve (pink) of a mixture of 6% *trans*- and 94% *cis*-OptoNAM-3 (corresponding to the 350 nm PSS determined by HPLC) assuming only the *trans* isomer is active. The 6% *trans* theoretical dose response curve is also superimposable to the one of OptoNAM-3 350 nm PSS. This shows that the activity of the 365 nm PSS and 350 nm PSS of OptoNAM-3 can entirely be explained by the amount of remaining *trans* isomer in the PSS.

**(C-D) OptoNAM-3 binds at the ifenprodil binding site. (C)** Dose-response curves of OptoNAM-3 on GluN1/GluN2B-ΔNTD receptors in the dark (black curve, $IC_{50}$ = 20 ± 3.9 µM, n = 3-5) or pre-illuminated with 365 nm (violet curve, $IC_{50}$ = 25 ± 6.1 µM, n = 3-5). OptoNAM-3 dose-response curves on wild type GluN1/GluN2B receptors are shown as dashed curves (black and violet for the dark and 365 nm PSS conditions, respectively). The remaining, low affinity inhibition of OptoNAM-3 on GluN1/GluN2B-ΔNTD receptors is likely due to a non-selective pore block of NMDARs at negative holding potentials, as previously shown for ifenprodil (7, 8). **(D)** Dose-response curves of *trans*-OptoNAM-3 in the dark in absence (black curve, $IC_{50}$ = 0.38 ± 0.03 µM, n= 5-21) and in presence of 0.2 µM ifenprodil (a concentration close to ifenprodil $IC_{50}$, orange curve). Ifenprodil increases OptoNAM-3 $IC_{50}$ by ~3-fold ($IC_{50}$ = 1 .0 ± 0.10 µM, n= 4-5 in presence of 0.2 µM ifenprodil), which is consistent with a competitive interaction between these two compounds.

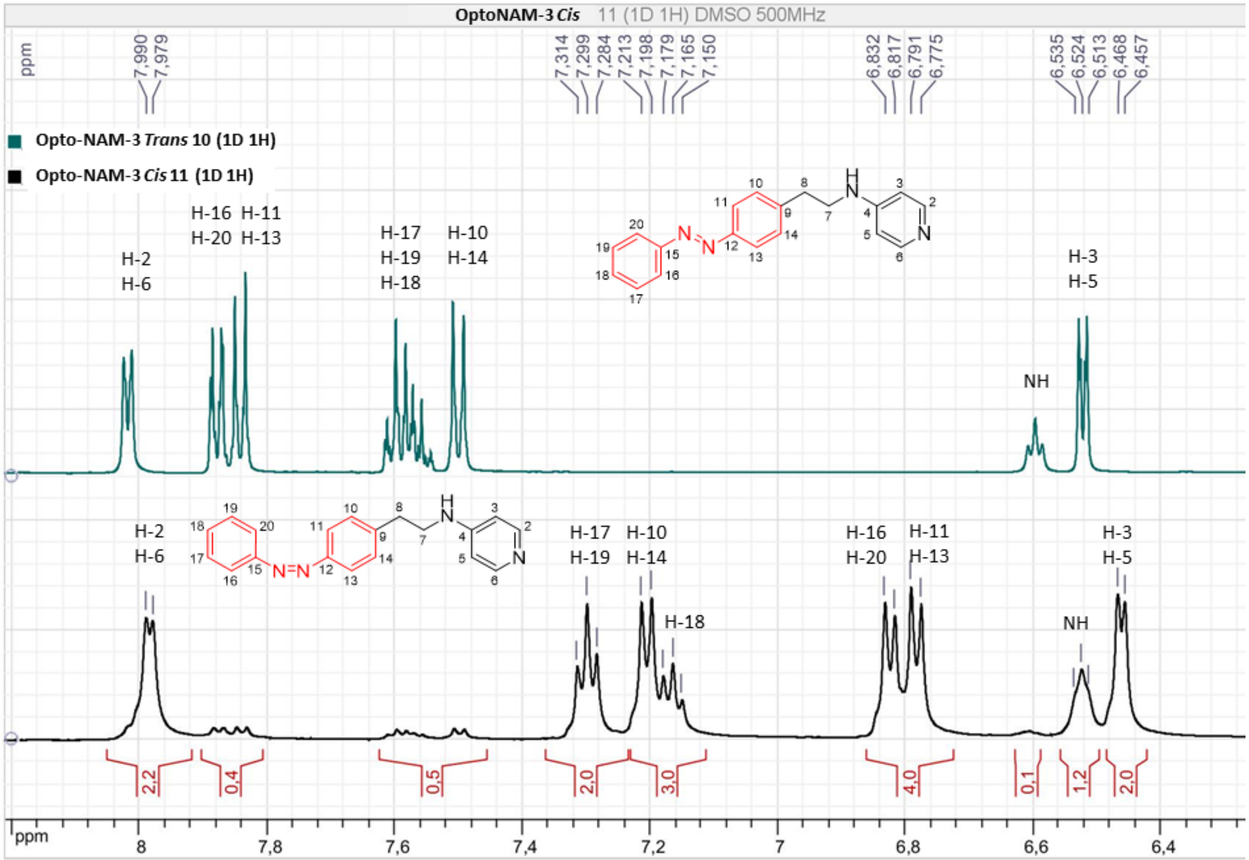


**Figure S4. ¹H NMR spectra of OptoNAM-3 dark and 365 nm PSS**. Upper panel: ¹H NMR spectrum of OptoNAM-3 in the dark, constituted of 100% *trans*. Lower panel: ¹H NMR spectrum of OptoNAM-3 365 nm PSS. The differences of chemical shifts of the hydrogen atoms allow separation of H atoms from the *cis* and the *trans* isomers. Percentage of *cis* and *trans* isomers PSS was calculated from the integration of the peaks at 6.8 ppm and 7.98 ppm.

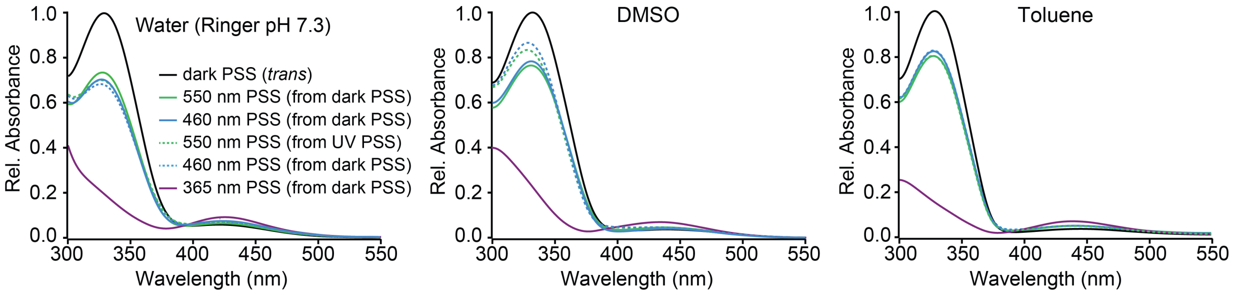


**Figure S5. OptoNAM-3 photochemical properties in different solvents.** UV-visible absorption spectra of OptoNAM-3 (25 µM) in water (Ringer at pH 7.3), DMSO and toluene. Black curves represent the dark PSS, the violet ones OptoNAM-3 365 nm PSS, the blue and green curves represent OptoNAM-3 460 nm and 550 nm PSS obtained after illumination of the dark PSS, and the dotted blue and green curves represent OptoNAM-3 460 nm and 550 nm PSS obtained after illumination of the 365 nm PSS.

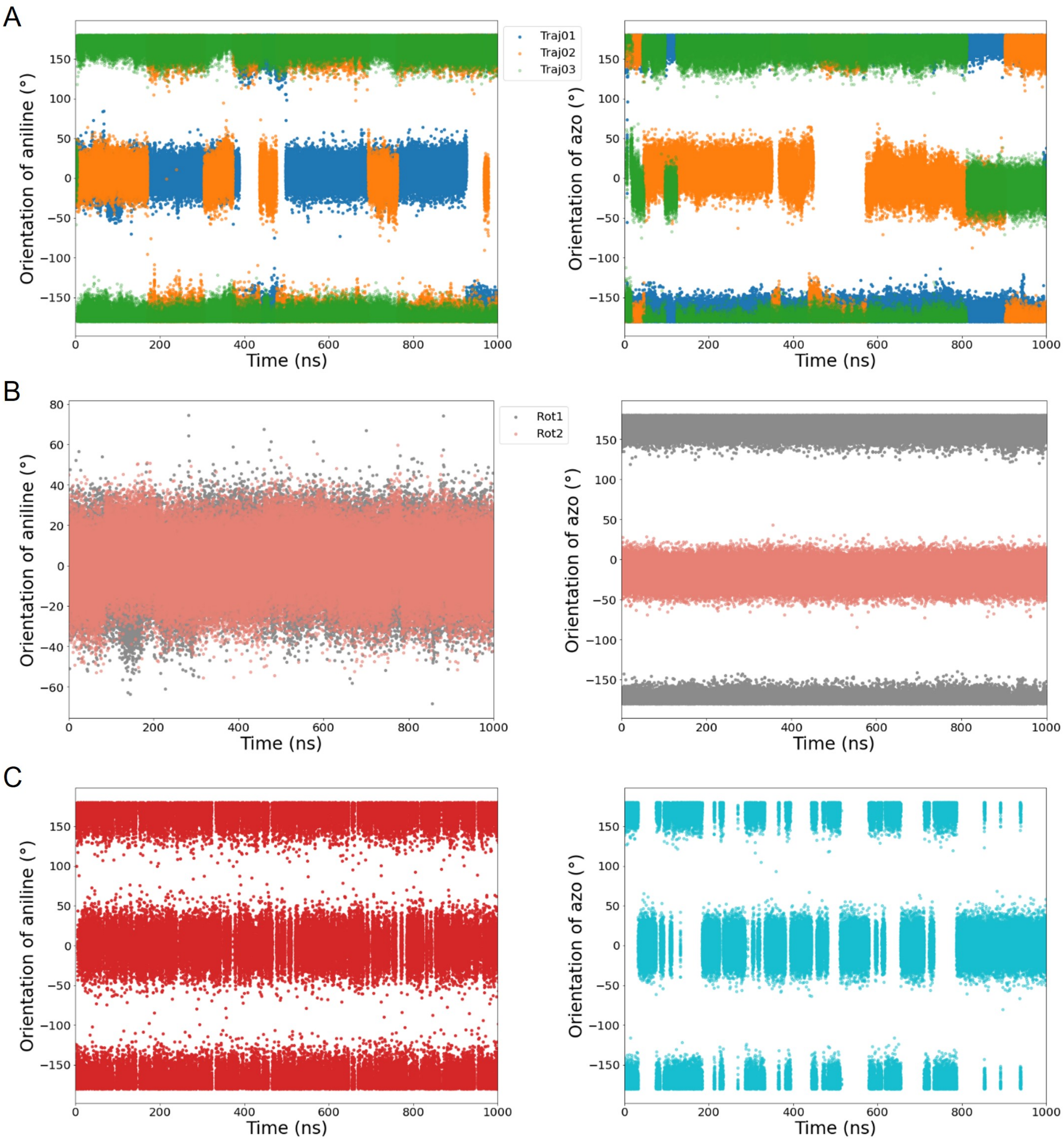
A
B
C
Traj01
Traj02
Traj03
Rot1
Rot2
Orientation of aniline (°)
Orientation of azo (°)
Time (ns)

**Figure S6. Additional data relative to Figure 5. (A)** Evolution of the two angles that describe OptoNAM-3 conformations. Left: orientation of the aniline. Right: orientation of the phenyl-azo moiety (C-C-N=N angle) (Fig. 5A). For the orientation of aniline, in trajectories 01 and 02 we observed exchanges between 0 and 180°, whereas in trajectory 03 the angle stayed at 180°. On average, this angle is 60.8% at 180° which means that this conformation is more stable by roughly 0.3 kcal/mol than the one at 0°. For the orientation of the azo moiety, in trajectory 01 the angle stayed at 180° and switched to 0° after 993 ns; for trajectory 02, we observed six conversions between the two basins; for trajectory 03, we observed some exchanges at the beginning, a stability from 125 to 810 ns, and then a final exchange. On average, this angle is 67.9% at 180° which means that this conformation is more stable by roughly 0.4 kcal/mol than the one at 0°. (**B**) Evolution of the same angles but for Rot1 and Rot2 OptoNAM-3 rotamers during the simulations under constraint. Left: orientation of the aniline. Right: orientation of the phenyl-azo moiety (C-C-N=N angle). **(C)** Evolution of the two angles for the ligand alone in water. For the orientation of aniline, the two basins at 0° and 180° are populated respectively 48.2% and 51.8% of the time, whereas for the orientation of the phenyl-azo they are populated at respectively 53.5% and 46.5% of the time. This corresponds to differences of free energies below 0.1 kcal/mol. We observed very low differences of free energies between the conformations of OptoNAM-3 both in the protein and in solution; however, in solution, we observed much more transitions between each conformation, which means that the free energy barrier to go from one conformation to the other is much smaller in solution than when bound to the protein.

**Figure S7.**

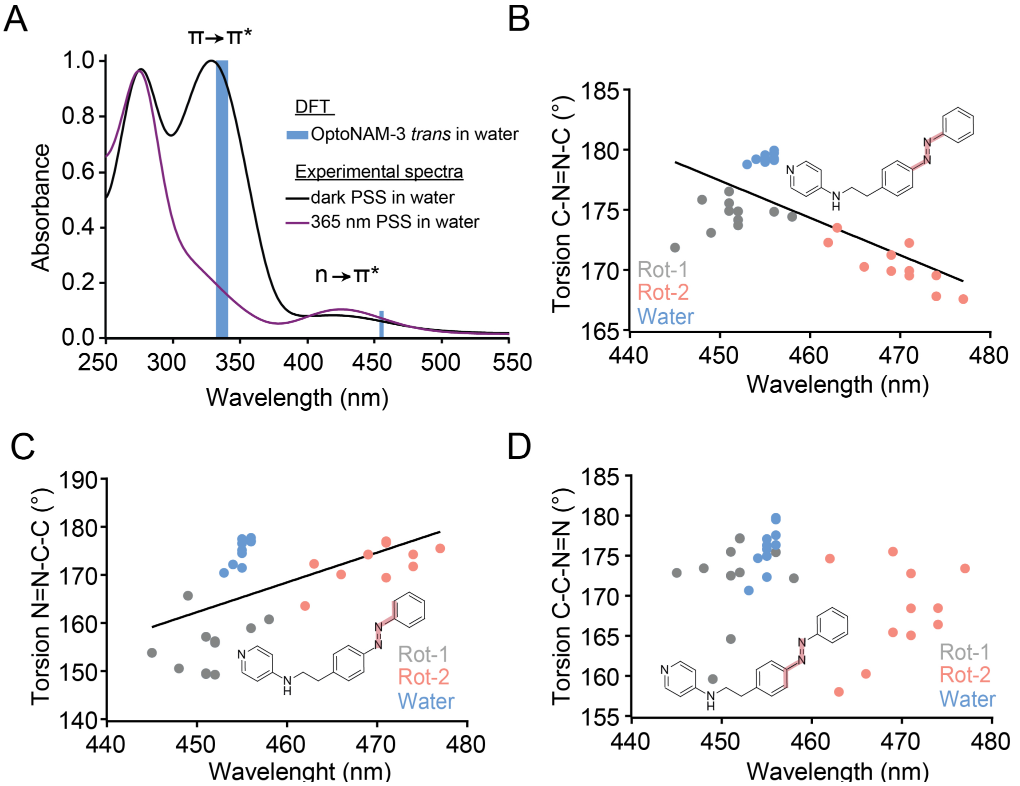


**Additional data relative to Figure 5. (A)** Superposition of the experimental π→ π* and n→π* bands of the UV-Vis spectra of OptoNAM-3 dark PSS (dark line) and 365 nm PSS (violet line), to the theoretical π→ π* and n→π* transitions of OptoNAM-3 in water predicted by DFT calculations (blue bars representing the range of predicted wavelengths for *trans*-OptoNAM-3 across the different snapshots of the dynamic, see Table S2). Relationships between the C-N=N-C **(B)**, N=N-C-C **(C)** and the C-C-N=N **(D)** torsion angles (as highlighted in pale red in the inset chemical structures) of OptoNAM-3 in the 11 snapshots selected for DFT calculations and their predicted n→π* absorption wavelengths, for Rot-1 (in grey), Rot-2 (in salmon) in their binding-site (as shown in Fig. 5D) and for *trans*-OptoNAM-3 in water (in blue). We can see in these representations that bound Rot-1 has a geometry much closer to OptoNAM-3 in water than Rot-2. In addition, geometry of *trans*-OptoNAM-3 in water is much more homogenous across the different snapshots of the dynamic (blues dots) compared to OptoNAM-3 Rot-1 and -2 in the protein (grey and salmon dots respectively).

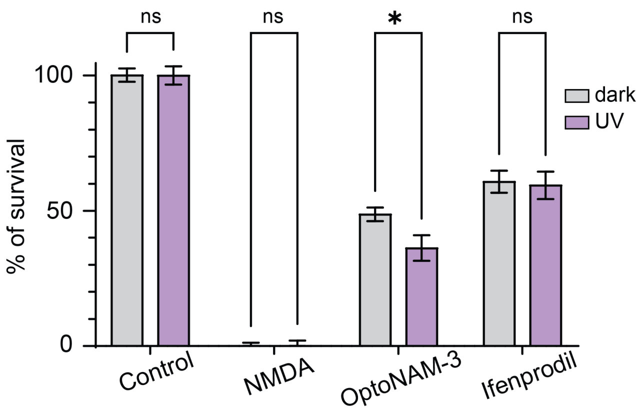


**Figure S8. OptoNAM-3 decreases NMDA-induced neuronal death in a photodependent manner.** Percentage of neuronal survival in cultured cortical neurons exposed either to control (0.01% DMSO and 10 µM Glycine), NMDA alone (100 µM NMDA and 10 µM Glycine), NMDA + ifenprodil or NMDA+ OptoNAM-3 (100 µM NMDA+ 5 µM of NAM), in the dark (grey bar) or after 2 min UV (365 nm) illumination (violet bar). In the presence of *trans*-OptoNAM-3 (dark) or ifenprodil, cell survival increased to 50% and 60%, respectively. When 365 nm illumination followed the addition of OptoNAM-3, cell survival was decreased to 35% but the extent of survival induced by ifenprodil was not affected, precluding any deleterious effect of the UV light treatment on cell survival. Multiple comparisons were performed by two-way ANOVA with Bonferroni's correction; n.s., $p > 0.05$; *, $p < 0.05$; n = 9-20 cultures per condition, in each culture 4-6 wells/condition. Data presented here (mean and SEM) were normalized to the control and NMDA conditions (see Methods for the calculation protocol).

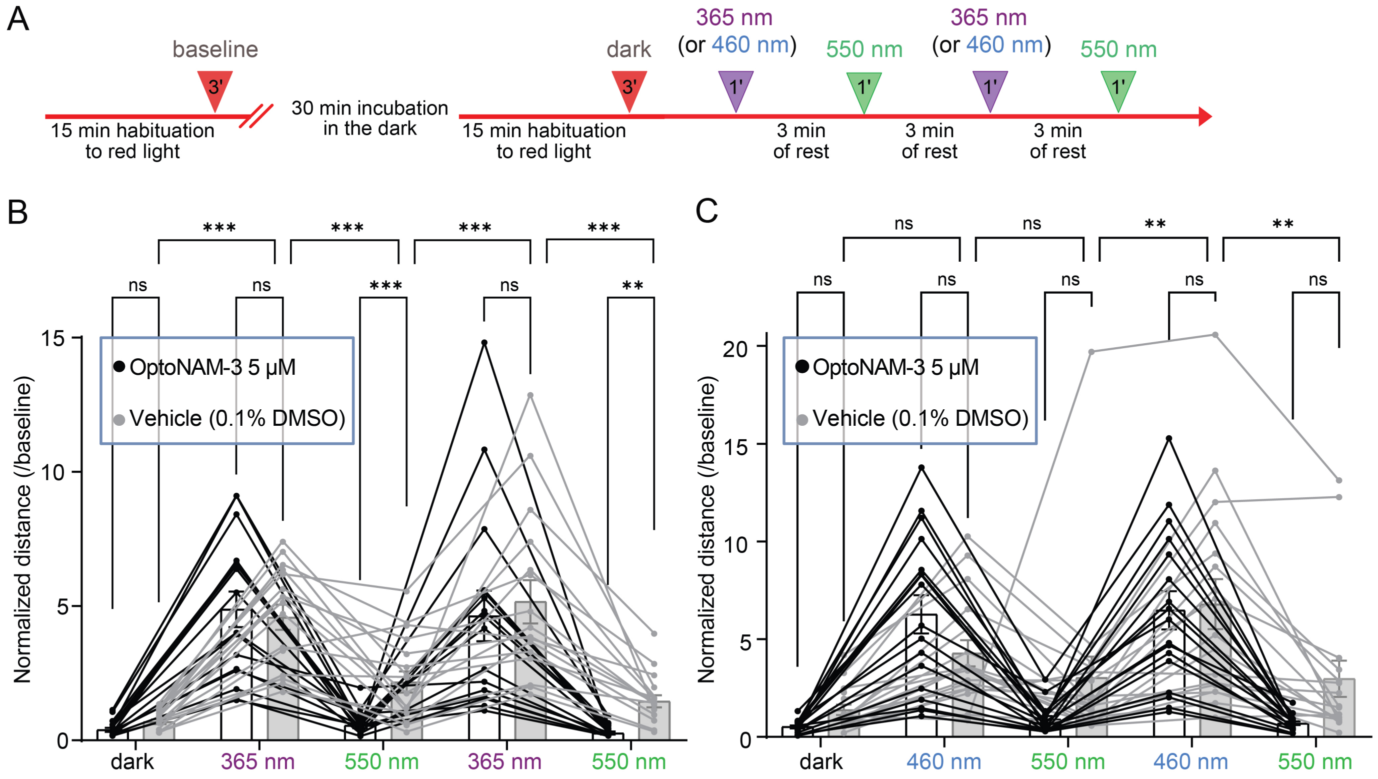


**Figure S9. OptoNAM-3 photomodulates Xenopus tadpole locomotion *in vivo*: protocol and tadpole locomotion normalized to baseline locomotion. Additional data relative to Figure 6. (A)** Experimental design of the behavioral tests performed on tadpoles incubated in control (0.1% DMSO) or OptoNAM-3 at 5µM and exposed to UV/green (or blue/green) light cycles. **(B, C)** Normalized distance (compared to baseline) traveled by tadpoles (1 point represents the mean distance traveled by the 3 tadpoles of one well) incubated in control (white bars and black points) or in 5 µM OptoNAM-3 (grey bars and points), in the dark and during UV/green light cycles (or blue-green light cycles for panel C). Note that UV light influences tadpole locomotion on its own. n = 16 wells for (B) and n = 17 wells for (C), which corresponds to a total of 48 and 51 tadpoles, respectively. The means of 3 tadpoles per well were used to conduct a paired statistical test. n.s., $p > 0.05$; ***, $p < 0.001$; **, $p < 0.01$; Friedman test followed by Dunn's multiple comparison test.

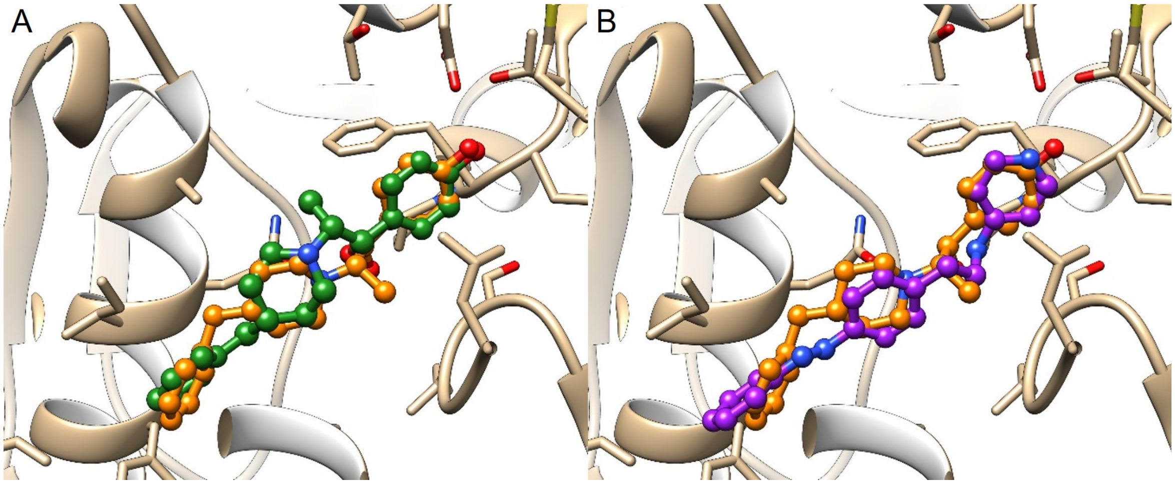


**Figure S10. (A)** Overlap between crystallographic (in orange) and docked (in green) poses of ifenprodil. **(B)** Overlap between the crystallographic pose of ifenprodil (in orange) and the docked pose of *trans*-OptoNAM-3 (in purple).

**Table S1.** Summary of the $IC_{50}$s of OptoNAMs in the dark and UV compared to the activity of their parent compounds (2–4, 9).

| OptoNAM- | Parent compound $IC_{50}$ or Ki (μM) | $IC_{50}$ (μM) In the dark (n) | $IC_{50}$ (μM) of the 365 nm [or 350 nm] PSS (n) | UV/dark |
|---|---|---|---|---|
| 1 | 0.003 [(a)] | 11 ± 1 (n= 4-6) | 37 ± 2 (n= 5-17) | 3.4 |
| 2 | 0.010 [(b)] | 24 ± 2 (n= 3-5) | 117 ± 10 (n= 3-8) | 4.9 |
| 3 | 0.093 [(c)] | 0.38 ± 0.03 (n= 5-21) | 1.7 ± 0.2 (n= 5-17)<br>[4.4 ± 0.6 (n= 4-13)] | 4.5<br>(11.5) |
| 4 | 0.00068 [(b)] | 0.024 ± 0.026 (n= 3) | 0.031 ± 0.029 (n= 3) | 1.3 |

(a) Ki from ref (3)
(b) $IC_{50}$ from ref (2)
(c) Ki from ref (9)
(d) Ki from ref (4)

**Table S2.** Computed vertical energies and oscillator strength of 11 snapshots of OptoNAM-3 in implicit water for the 2 first visible transitions using the B2PLYP functional.

| Snapshot (MD time) | Computed vertical energy in eV (nm) and oscillator strength | | Snapshot (MD time) | Computed vertical energy in eV (nm) and oscillator strength | |
|---|---|---|---|---|---|
| | Transition 1 (n → π*) | Transition 2 (π → π*) | | Transition 1 (n → π*) | Transition 2 (π → π*) |
| 0 (3 μs) | 2.72 (456) f = 0.000029 | 3.64 (341) f = 0.884 | 6 (3.6 μs) | 2.72 (455) f = 0.0038 | 3.65 (339) f = 0.917 |
| 1 (3.1 μs) | 2.72 (456) f = 0.00023 | 3.69 (336) f = 1.093 | 7 (3.7 μs) | 2.72 (455) f = 0.0021 | 3.69 (335) f = 1.09 |
| 2 (3.2 μs) | 2.73 (454) f = 0.0067 | 3.69 (336) f = 1.05 | 8 (3.8 μs) | 2.73 (455) f = 0.0093 | 3.72 (339) f = 1.07 |
| 3 (3.3 μs) | 2.73 (455) f = 0.0042 | 3.70 (335) f = 1.08 | 9 (3.9 μs) | 2.72 (456) f = 0.00062 | 3.69 (336) f = 1.09 |
| 4 (3.4 μs) | 2.74 (453) f = 0.013 | 3.73 (332) f = 1.06 | 10 (4 μs) | 2.73 (455) f = 0.0021 | 3.67 (338) f = 1.12 |
| 5 (3.5 μs) | 2.72 (456) f = 0.00065 | 3.65 (340) f = 0.92 | | | |

**Table S3:** Computed vertical energies and oscillator strength of 11 snapshots of OptoNAM-3 inside the protein for the 2 rotamers and for the 1st transition (n → π*) using the B2PLYP functional.

| Snapshot (MD time) | Computed vertical energy in eV (nm) and oscillator strength | | Snapshot (MD time) | Computed vertical energy in eV (nm) and oscillator strength | |
|---|---|---|---|---|---|
| | Rot1 (n → π*) | Rot2 (n → π*) | | Rot1 (n → π*) | Rot2 (n → π*) |
| 0 (900 ns) | 2.74 (452)<br>f = 0.00073 | 2.64 (469)<br>f = 0.038 | 6 (960 ns) | 2.70 (458)<br>f=0.00002 | 2.68 (462)<br>f = 0.0089 |
| 1 (910 ns) | 2.77 (448)<br>f=0.0019 | 2.63 (471)<br>f = 0.021 | 7 (970 ns) | 2.72 (456)<br>f=0.0013 | 2.63 (471)<br>f = 0.037 |
| 2 (920 ns) | 2.74 (452)<br>f=0.0021 | 2.61 (474)<br>f=0.045 | 8 (980 ns) | 2.76 (449)<br>f=0.0008 | 2.66 (466)<br>f=0.044 |
| 3 (930 ns) | 2.74 (452)<br>f=0.0029 | 2.61 (474)<br>f=0.036 | 9 (990 ns) | 2.75 (451)<br>f=0.0016 | 2.68 (463)<br>f=0.029 |
| 4 (940 ns) | 2.79 (445)<br>f=0.00009 | 2.64 (469)<br>f = 0.015 | 10 (1000 ns) | 2.75 (451)<br>f=0.00629 | 2.63 (471)<br>f=0.039 |
| 5 (950 ns) | 2.75 (451)<br>f=0.00031 | 2.60 (477)<br>f = 0.038 | | | |

**Table S4:** Computed vertical energies and oscillator strength of snapshot 0 of OptoNAM-3 for the 2 rotamers and for the 1st transition (n → π*), inside the protein (**1**, first line); without the protein without optimization (**2**, second line), and without the protein after optimized in vacuum (**3**, third line)

| Snapshot (MD time) | Computed vertical energy in eV (nm) and oscillator strength | | |
|---|---|---|---|
| | Rot1 (n → π*) | Rot2 (n → π*) | |
| 0 (900 ns) | 2.74 (452)<br>f = 0.00073 | 2.64 (469)<br>f = 0.038 | **1** |
| 0 (900ns) without protein, in vacuum (electrostatic contribution) | 2.72 (456)<br>f = 0.0012 | 2.63 (471)<br>f = 0.037 | **2** |
| 0 (900ns) without protein + optimization in vacuum (geometric contribution) | 2.65 (467)<br>f = 0.0011 | 2.65 (467)<br>f = 0.000046 | **3** |

### SI References


1. E. Merino, M. Ribagorda, Control over molecular motion using the cis-trans photoisomerization of the azo group. *Beilstein J Org Chem* **8**, 1071–1090 (2012).

2. J. A. McCauley, NR2B subtype-selective NMDA receptor antagonists: 2001 – 2004. *Expert Opinion on Therapeutic Patents* **15**, 389–407 (2005).

3. B. Büttelmann, *et al.*, 4-(3,4-dihydro-1H-isoquinolin-2yl)-pyridines and 4-(3,4-dihydro-1H-isoquinolin-2-yl)-quinolines as potent NR1/2B subtype selective NMDA receptor antagonists. *Bioorg Med Chem Lett* **13**, 1759–1762 (2003).

4. J. A. McCauley, *et al.*, NR2B-SelectiveN-Methyl-D-aspartate Antagonists: Synthesis and Evaluation of5-Substituted Benzimidazoles. *J. Med. Chem.* **47**, 2089–2096 (2004).

5. I. Kaljurand, T. Rodima, I. Leito, I. A. Koppel, R. Schwesinger, Self-consistent spectrophotometric basicity scale in acetonitrile covering the range between pyridine and DBU. *J Org Chem* **65**, 6202–6208 (2000).

6. B. Büttelmann, *et al.*, 4-(3,4-Dihydro-1H-isoquinolin-2yl)-pyridines and 4-(3,4-Dihydro-1H-isoquinolin-2-yl)-quinolines as potent NR1/2B subtype selective NMDA receptor antagonists. *Bioorganic & Medicinal Chemistry Letters* **13**, 1759–1762 (2003).

7. F. Perin-Dureau, J. Rachline, J. Neyton, P. Paoletti, Mapping the Binding Site of the Neuroprotectant Ifenprodil on NMDA Receptors. *J. Neurosci.* **22**, 5955–5965 (2002).

8. L. Mony, *et al.*, Structural basis of NR2B-selective antagonist recognition by N-methyl-D-aspartate receptors. *Mol Pharmacol* **75**, 60–74 (2009).

9. N. J. Liverton, *et al.*, Identification and characterization of 4-methylbenzyl 4-[(pyrimidin-2-ylamino)methyl]piperidine-1-carboxylate, an orally bioavailable, brain penetrant NR2B selective N-methyl-D-aspartate receptor antagonist. *J Med Chem* **50**, 807–819 (2007).
